# Delayed viral vector mediated delivery of neurotrophin-3 improves skilled hindlimb function and stability after thoracic contusion in rats

**DOI:** 10.1101/2022.02.11.479866

**Authors:** Jared D. Sydney-Smith, Alice M. Koltchev, Lawrence D. F. Moon, Philippa M. Warren

## Abstract

It has been reported that intramuscular injection of an Adeno-associated viral vector serotype 1 (AAV1) encoding Neurotrophin-3 (NT3) into hindlimb muscles 24 hours after a severe T9 contusion in rats induced lumbar spinal neuroplasticity, partially restored locomotive function and reduced spasms during swimming. Here we investigated whether a targeted delivery of NT3 to lumbar and thoracic motor neurons 48 hours following a severe contusive injury aids locomotive recovery in rats. AAV1-NT3 was injected into the *tibialis anterior, gastrocnemius* and *rectus abdominus* muscles 48-hours following trauma, persistently elevating serum levels of the neurotrophin. NT3 improved trunk stability, accuracy of stepping during skilled locomotive tasks, and alternation of the hindlimbs during swimming, but it had no effect on gross locomotion function in the open field. The number of vGlut1^+^ (likely proprioceptive afferent) boutons on *gastrocnemius* α-motor neurons was increased after injury but normalised following NT3 treatment suggestive of a mechanism in which the functional effects may be mediated through proprioceptive feedback. *Ex vivo* MRI revealed substantial loss of grey and white matter at the lesion epicentre but no effect of delayed NT3 treatment to induce neuroprotection or prevent secondary damage. Spasms and hyperreflexia were not reliably induced in this severe injury model suggesting a more complex anatomical or physiological cause to their induction. We have shown that delayed intramuscular AAV-NT3 treatment can promote recovery in skilled stepping and coordinated swimming supporting a role for NT3 as a therapeutic strategy for spinal injuries potentially through modulation of somatosensory feedback.

**Key Points:**

1. Targeted delivery of NT3 to hindlimb and trunk muscles at a clinically relevant 48h following a severe thoracic contusion aids fine locomotor control and synchronised movement.
2. NT3 mediated improvements in trunk stability, accuracy of stepping during skilled locomotive tasks, and alternation of the hindlimbs during swimming through the normalisation of vGlut1+ boutons on presumptive proprioceptive afferents innervating these muscles.
3. 250kDyn thoracic contusion does not reliably result in measurable signs of spasticity.

## Introduction

Most traumatic spinal cord injuries (SCIs) are incomplete, leaving patients with some degree of residual sensory and motor function below the neurological level of the injury (National Spinal Cord Injury Statistical Centre. 2019) while causing life-altering deficits including limb paralysis and spasticity. 25-55% of SCIs occur within the thoracic or lumbar spinal cord, causing reduced lower limb function and trunk instability (Divanoglou and Levi, 2009, Pickett et al., 2003, Singh et al., 2014). This is due to damage to the descending supraspinal and propriospinal axons which project below the lesion and are critical for specific functions, such as fine locomotor control and synchronised movements (Jordan et al., 2008).

Some degree of spontaneous motor recovery occurs in both human patients and pre-clinical SCI models following incomplete injuries due to the formation of novel circuits which bypass the injury site to reinnervate circuits below the lesion and establish basic motor function (Nishimura et al., 2007, Zorner et al., 2014, Zörner et al., 2010, Tohyama et al., 2017). Proprioceptive feedback involving vGlut1+ afferents is essential for this motor recovery (Edgerton et al., 2008, Rossignol and Frigon, 2011, Takeoka et al., 2014), enabling information related to movement to reach the spinal cord circuits, stabilising and refining motor output. Nonetheless, use of any residual function and attempts to further recovery through rehabilitation can be disrupted by the development of spasticity which occurs in up to three quarters of patients (Maynard et al., 1990, Skold et al., 1999), and results in involuntary spasms, rigidity and exaggerated reflexes (Mukherjee and Chakravarty, 2010, Pandyan et al., 2005). As such, it is important to establish reliable strategies to enhance growth and circuit reformation from the descending axons to motoneurons within the lumbar spinal cord.

Neurotrophic factors support the activity and function of descending spared axons following SCI (da Silva and Wang, 2011). Neurotrophin-3 (NT3) is a growth factor required for correct connectivity between proprioceptive afferents from muscle to the spinal cord (Boyce and Mendell, 2014b, Boyce and Mendell, 2014a). We, and others, have provided supplementary NT3 in animal models of neurological injury and shown that it enhances neuroplasticity and improves sensorimotor recovery in mice, rats and cats (Kakanos and Moon, 2019, Boyce and Mendell, 2014b, Boyce and Mendell, 2014a, Fortun et al., 2009, Krupka et al., 2017, Ollivier-Lanvin et al., 2015, Sydney-Smith et al., 2021a, Han et al., 2019), while clinically it has been shown not to cause pain (Parkman et al., 2003, Coulie et al., 2000, Chaudhry et al., 2000, Sahenk, 2007, Sahenk et al., 2014). Delivery of NT3 via injection of AAV1 into forelimb flexor muscles reversed some sensorimotor deficits after bilateral corticospinal tract lesion in the brainstem of rats (Kathe et al., 2016b) and improved functional outcomes after mid cervical contusion injury in rats (Sydney-Smith et al., 2021a). Similarly, a recent study assessed whether AAV-NT3 induced therapeutic effects when injected into hindlimb muscles 24-hours after a T8 thoracic contusion (Chang et al., 2019). NT3 treatment reduced the presence of hindlimb spasms during swimming and normalised hyperreflexia in the hind paw. The study also revealed that hindlimb rehabilitation reduced hyperreflexia and the number of spastic events during swimming by six weeks post injury. There was however no synergistic effect of rehabilitation and NT3 treatment on functional and electrophysiological outcomes.

We hypothesised that intramuscular injection of AAV1-NT3 into muscles innervated by thoracic and lumbar spinal motor neurons would enhance skilled motor function and potentially alleviate effects of hindlimb hyperreflexia and spasticity if delivered at a clinically relevant 48-hours following the initial trauma. We show that severe thoracic contusion led to persistent hindlimb deficits but few, if any, spasms or hyper-reflexivity. Nevertheless, 48h-delayed NT3 treatment improved skilled locomotor function and hindlimb alternation during swimming potentially due to modifications of vGlut1 signalling on proprioceptive afferents.

## Methods

### Ethical approval and animal welfare

Experiments were approved by the King’s College London Welfare and Ethics Committee and were conducted in accordance with UK Animals (Scientific Procedures) Act 1986 (ASPA) under Home Office Project Licence number P53631BC2. They have been performed in accordance with the ethical standards laid down in the 1964 Declaration of Helsinki and its later amendments. During all experiments, data processing, and analysis investigators were blind as to the treatment group of each animal. An independent third party coded and randomly assigned animals to treatment groups (simple randomisation) without knowledge of injury impact based on behavioural assessment or electrophysiological recordings.

Animals were housed in groups of four or five, exposed to a normal 12-hour dark-light cycle at 21C with access to food, water and environmental enrichment *ad libitum*. The health and welfare of all animals was monitored daily by veterinary staff and the study investigators at King’s College London and was in accordance with the Animal Welfare Act 2006. 28 female Wistar rats (256g ± 29g; Envigo) were used in this study. Of the 28 animals, one died three days after contusion injury, with all other animals making uneventful recoveries. One animal was excluded from the study due to full locomotor recovery (assessed through a bilateral (BBB) score of 21) within one week of injury. Animals were divided into 3 groups: 1) contusion + NT3 (NT3, n=11); 2) contusion + PBS (PBS, n=12); and 3) uninjured + no treatment (sham, n=4). Following injury animals were housed in mixed cages comprising injured animals of both NT3 and PBS treated animals for the duration of the study with sham animals housed together in a separate cage.

### Injury and treatment application

#### Thoracic contusion injury

Rats were anaesthetised with Isoflurane (5%; Zoetis, UK Ltd) at 1 L.min-1 O_2_ flow (0.4 L.min-1 kg-1) induction, and maintained with 2.5% isoflurane (0.4 L.min-1 kg-1) throughout surgery. Depth of anaesthesia was assessed throughout surgery through continuous monitoring of the pedal reflex, respiration rate and pattern, and colour of mucous membranes. Preoperatively 5mg/kg Enrofloxacin (Baytril) and 5 mg/kg carprofen (Carprieve) were injected subcutaneously. Body temperature was maintained throughout the surgery at 37±1°C using a homothermic heat pad (Harvard Apparatus). Upon reaching a surgical plane of anaesthesia, eye ointment (Viscotears) was applied and the region around the lower thoracic vertebrae was shaved, the skin prepared with both 4% w/v chlorhexidine gluconate (Hibiscrub) and iodine antiseptic (Vetark Professionals).

Using sterile techniques and surgical antisepsis, the thoracic (T) spinal column was exposed through completion of a 2cm dorsal midline incision from T8-T11, and subsequent retraction of the skin and spinotrapezius muscles. While preserving the facet joints and dura, a laminectomy performed over T8 exposing the spinal cord. The vertebral column was clamped at the T8-T9 vertebra using the Infinite Horizon contusion impactor (Precision Systems and Instrumentation). Animals received a single bilateral contusion, of 250 kDyn force with 0s dwell time using a 2.5mm diameter impact tip (Precision Systems Instrumentation) at the T9 spinal segment. Injury completeness was confirmed through *ex vivo* MRI analysis. Muscle layers and the skin were sutured in two layers (4-0 Vicryl; Ethicon) and animals were given saline (5mL; s.c.) and recovered in a heated environment (30°C; Thermocare) before transfer to their home cage. Sham animals received a full T8 laminectomy, without clamping in the Infinite Horizons impactor, and all pre- and post-operative drugs.

For 5 days after injury animals continued to receive; warm saline (s.c.), 5 mg/kg Baytril, and 5mg/kg Carprieve in addition to nutritional support if the animal’s weight fell 5% below that determined pre-injury and were placed in incubator cages maintained at 24-25°C. Animals were unable to void their bladders naturally for the first week after injury, necessitating manual bladder emptying three times per day. Once spontaneous voiding was recovered, bladder volume and voiding was monitored daily to ensure no infections occurred.

### Adeno-associated virus encoding neurotrophin-3

An AAV transfer plasmid, AAVspNT3, was cloned to contain the full length human prepro-neurotrophin-3 under a CMV promoter. The neurotrophin-3 coding DNA sequence (NM_002527.4), corresponding to isoform 2 precursor protein including the secretory signal, was flanked by splice donor/splice acceptor sites and terminated in a beta globin poly(A) sequence (gifted by Prof. Fred Gage, Salk Institute). This was packaged into an AAV serotype 1 (University of Pennsylvania Viral Vector Core facility; custom batch CS1232). This vector (referred to as AAV-NT3) was titred using digital droplet PCR (6.2 x10^13^ genomic copies (GC)/ml; Viral Vector Core) and in our lab via qPCR (1.22 x10^13^ GC/ml), the latter value being used for dilution calculations. Primers for the qPCR were: Forward 5’-AATTACCAGAGCACCCTGCC and Reverse 5’-TTTTGATCTCCCCCAGCACC.

### Intramuscular injections

Forty-eight hours after contusion, the *tibialis anterior* and *gastrocnemius* muscles of both hindlimbs as well as *rectus abdominus* were injected with AAV-NT3 or PBS (Figure 1A). Similar to the injury surgery, using sterile techniques and surgical antisepsis, the animals were anaesthetised (isoflurane), given eye ointment, Enrofloxacin, and Carprieve. Both hindlimbs and the abdomen were shaved, and the skin sterilised. Following each set of muscle injections, the incision was sutured (4-0 Vicryl). Injections were made using a 26G non-coring bevel needle attached to a Hamilton syringe. A total of 2.57 x10^12^ genomic copies of AAV-NT3 diluted in 220μl of PBS (or equivalent amount of PBS vehicle) was injected into each animal. Injections into the *tibialis anterior* and *gastrocnemius* muscles targeted motor end plates deep within the hindlimb muscles (Yin et al., 2019) and muscle spindles (Figure 1B). For the *rectus abdominus* all injections were superficial, with care taken not to penetrate through the muscle into the peritoneal cavity.

**Figure 1:**
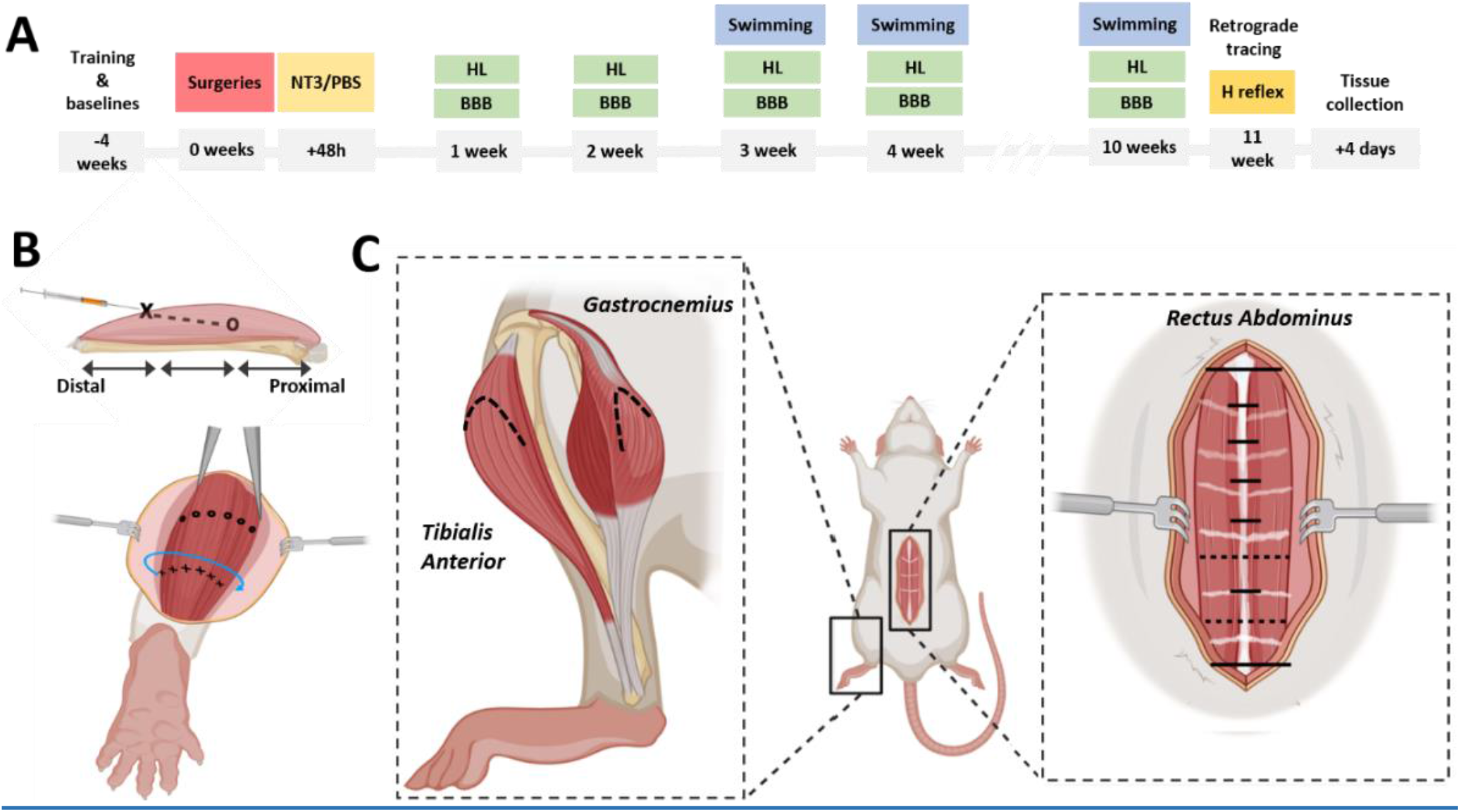
Experimental overview. A) Timeline indicating the timepoints of each behavioural and electrophysiological assay. BBB= Basso, Beattie and Bresnahan locomotor scale, and HL=horizontal ladder. B) Top panel: cross section through a muscle indicating the approximate location of needle insertion (X) and injection site location and depth (O). Hindlimb muscles were injected distoproximally as shown. Bottom panel: circumferential arc around the muscle belly (blue), with points of insertion shown, designed to target the entirety of the muscle belly. C) Schematic of the rat in a supine position showing the injected muscles. Left: lateral hindlimb showing the *tibialis anterior* and *gastrocnemius,* with medial head in front and paler compared to the lateral. Right: *rectus abdominus*, showing the tendinous insertions (pale pink) and the nine imaginary transverse lines (black lines) used to approximate the injection sites (black dashed lines). Long solid black lines indicate the xyphoid process and the pubic symphysis, dashed lines indicate the approximate positions of the injections.

#### Tibialis Anterior

An incision was made on the anterolateral aspect of the lower hindlimb. Injections were made across the proximal aspect of the muscle from the ankle to the knee. A total of 6×5μl of AAV-NT3 (3.50×10^11^ genomic copies) or PBS was made into each *tibialis anterior*, with final injection sites running in a steep circumferential arc across the entire fascial compartment (Figure 1C).

#### Gastrocnemius

An incision was made along the posterior aspect of the lower hindlimb extending one third the distance proximally from the popliteal fossa to the ankle. A total of 6×5μl of AAV-NT3 (7.1×10^11^ genomic copies) or PBS was made into both the medial and lateral heads of the *gastrocnemius*, with the final injection sites positioned in a steep circumferential arc across the thickest part of the muscle belly.

#### Rectus Abdominus

An incision was made from the xiphoid process down to the pubic symphysis. Injections were made between the 4^th^-6^th^ tendinous insertions as this portion of the muscle is innervated at or below the level of thoracic contusion (Hijikata et al., 1992) (Figure 1C). Two 5μl injections of AAV-NT3 (4.68 x10^11^ genomic copies) or PBS were bilaterally delivered equating to 20μl total delivered to each side of the *rectus abdominus*.

### Behavioural functional assessment

#### Open field locomotor assessments

Locomotor functional recovery was monitored using the 0-21 point Basso, Beattie and Bresnahan scale (BBB) (Basso et al., 1995). 1 week prior to injury, animals were habituated to a circular Perspex open field assessment area (100cm diameter, 20cm high) for ten minutes each day. Testing was done in the afternoon at day −1, and then weekly following injury during a 5 min period by 2 assessors in a fluorescently-lit room.

#### Analysis

The BBB score for each hindlimb were averaged between the two assessors and the BBB sub-scores similarly calculated (Popovich et al., 2012).

### Horizontal ladder

Accuracy of hindlimb placement was assessed through the animals’ ability to cross a 1m horizontal ladder with rungs spaced randomly 1-3cm apart. Prior to injury, animals were trained to cross the ladder for 15 min periods each day and baseline values obtained on day −1 prior to injury. Testing was done in the afternoon weekly following injury in a fluorescently-lit room. 3-5 complete runs across the ladder were video recorded.

#### Analysis

For analysis, steps on both hindlimbs were assessed using seven-point horizontal ladder scoring system (Metz and Whishaw, 2009). Videos were analysed in slow motion using VLC Media Player (VideoLan).

### Swimming

To assess non-weight bearing locomotor function, animals had to swim in a rectangular glass chamber (120cm x 12cm x 50cm) filled with 25cm water at 20-23°C (Ryu et al., 2017, Gonzenbach et al., 2010). A Perspex mirror was placed at a 45° angle on the base of the swimming tank to record a dorsal and lateral views of the animal. Rats placed at one end of the chamber would swim to an accessible rigid platform (Figure 5A) at the other end. Animals were habituated to the chamber daily for 2 weeks prior to injury and baseline recordings collected on day −1. Animals were assessed weekly from 3 weeks following injury. Animals were recorded (HERO7 Black, GoPro) over an 80cm distance (= 1 run). 5-7 complete runs were filmed before the animal was towel dried and returned to their home cage. The testing was carried out at the same time of day.

#### Analysis

Over 5 videotaped runs, hind-limb swim strokes during normal ‘left-right alternating’ swimming during slow motion. The percentage of left-right alternating and total number of strokes was calculated for each limb. Hindlimb strokes were excluded if they: occurred outside the 80cm assessment area; were associated with a change in direction; or they were the initial stroke following an interruption to swimming. The number of coordinated hind limb strokes over all the runs performed by each animal was displayed as a proportion of total number of hindlimb strokes performed.

### H reflex recordings

Animals were anaesthetised with ketamine and medetomidine (respectively 30mg/kg and 0.10 mg/kg i.p.) and were treated with atropine sulphate. Depth of anaesthesia was assessed as previously described. Preoperatively 5 mg/kg carprofen (Carprieve) was injected subcutaneously. Body temperature was maintained at 37±1°C using a homothermic heat pad (Harvard Apparatus). Upon reaching a surgical plane of anaesthesia, eye ointment (Viscotears) was applied and the ankle and hindpaw cleaned with 4% chlorhexidine. Two 26-gauge needles were positioned, subcutaneously in parallel and approximately 2mm apart, over the medial malleolus of the ankle to stimulate the medial plantar nerve. To record compound muscle action potentials (CMAPs) from the intrinsic hindpaw muscles innervated by the medial plantar nerve, two non-insulated needle recording electrodes were positioned at the base of first digit between the two prominent paw pads and placed superficially, 1mm apart and 2mm deep.

A monophasic square wave stimulus (width 100μs) was applied using an isolated constant current pulse stimulator (DS3, Digitimer) and the CMAP signal (amplified 2000×, band-pass filtered between 300Hz - 6kHz) was recorded and digitized via a Power1401-3A unit (Cambridge Electrical Design Ltd, CED) and visualised in Signal software (CED), recording in 5000ms sweeps. Motor threshold was again defined as the stimulus at which the M wave was elicited in three out of four stimuli. A recruitment curve was generated by recording 15 stimuli at 0.2HZ, starting at 1x motor threshold (MT) to a maximum of 2x MT in 10% increments up to 1.5xMT and finally at 2xMT. Subsequent rate dependent depression (RDD) was assessed with a paired stimulation protocol (Figure 8) at 1.3xMT stimulus intensity. Every 5000ms a pair of pulses were delivered. The first pulse occurred 300ms into the 5000ms recording sweep. Fifteen paired pulses were delivered at each of the following interstimulus intervals in this specific order: 100ms, 500ms, 50ms, 1000ms, 20ms, 700ms, 10ms, 20ms, 2000ms and 5000ms. Once recordings were finished, 2mg/kg (i.p.) atipamezole hydrochloride and 5mL saline (s.c.) was applied. Animals were assessed at the experimental end-point.

#### Analysis

Amplitude of the M and H waves was calculated in Signal software. The recruitment curve yielded the maximum H wave amplitude (Hmax) and maximum M wave amplitude (Mmax). For each interstimulus interval, the 15x M wave and H wave amplitudes corresponding to the conditional stimulus and the 15 for the test stimulus were pooled and averaged. These averages were then normalised to the Mmax for that testing session. Finally, RDD was calculated by representing the normalised H wave of the test stimulus as a percentage of the H wave amplitude with the conditioning stimulus.

### Retrograde tracing

Cholera Toxin Subunit B (CTb, #103, List Biologicals) was used to retrogradely trace motor neurons innervating the lateral head of the *gastrocnemius* in the left hindlimb. Tracing occurred immediately following H reflex recordings. The anaesthetised animals hindlimbs were shaved, the skin sterilised, and the muscle incised as described previously. Using a 26G needle on a Hamilton syringe, 5×1μl of 0.5% Cholera Toxin B (diluted in ddH_2_0) was injected into the lateral muscle head. The skin was sutured as described previously. Following surgery, animals were given 5ml of saline and the following day a single dose of 5mg/kg s.c. Carprieve.

### Tissue collection

Animals were deeply anaesthetized with sodium pentobarbital (Euthatal, 80 mg/kg i.p.). 1.5ml of blood was taken through cardiac puncture of the left ventricle and stored overnight at 4°C. Animals were transcardially perfused with PBS. The left *gastrocnemius*, including both the medial and lateral heads, was removed and snap frozen at stored at −80°C. Animals were perfused with 4% paraformaldehyde in 0.1M phosphate buffer. The spinal cord from C3 to the cauda equina was removed and post fixed in fresh 4% PFA in PBS for 24 h at 4°C.

### ELISA & BCA quantification

Levels of NT3 protein in the serum were assessed using a Human Neurotrophin-3 ELISA Kit (#ab100615 Abcam). As the viral vector encoding the human prepro-NT3 gene is identical to mature rat NT3, all NT3 (viral and endogenous) is detected. The lower limit of detection is 4.12pg/ml. Whole blood was centrifuged (7200g, 25°C) and the serum removed. The serum was centrifuged again, aliquoted and stored at −80°C. ELISA was performed as per the manufacturer’s instructions. The resulting plate was read immediately at 450nm using a BMG LabTech FLUOStar (Omega). NT3 levels were normalised to the total amount of protein extracted from each sample. A Bicinchoninic acid assay (BCA) assay (#71285-3, Novagen, Millipore) was additionally performed according to manufacturer’s protocol. Serum samples were diluted 1:150 in PBS, and a four parameter standard curve was generated (above r>0.99).

### Ex vivo *MRI*

For lesion quantification the epicentre of the lesion was imaged using a 9.4 T MRI scanner (Brucker Biospec) to acquire T2 weighted images. This was acquired using a fast spin-echo sequence: echo train length = 4, effective TE = 38 ms, TR = 3000 ms, FOV = 40 x 20 x 20 mm, acquisition matrix = 400 x 200 x 200, acquisition time = 9 h 20m. This composite scan contained T2W images from 14 spinal cords scanned simultaneously. The spinal cord was imaged from T6 to L1. 14 cords were imaged using a custom-made immobilising device. The tissue was fully submerged in Fomblin (Solvay) and all air bubbles were removed prior to scanning. The spinal cords were prepared for scanning as described previously (Sydney-Smith et al., 2021a) with additional care taken due to the reduced stability of the spinal cord around the epicentre compared to C5-C6 contusion.

#### Analysis

ITK -snap and Convert3D software were used to process the T2 weighted images. Convert 3D was used to alter the voxel dimensions to isotropic voxel size of 50μm. The composite image was separated into individual spinal cords to final voxel dimensions of 30×100×30μm in the *X* x *Y* x *Z* axis respectively. Within ITK-snap the contrast was increased (minimum adjustment:0.03, maximum adjustment:0.82, levels:0.41, window:0.82). Automatic segmentation was performed to select: presumptive spared white matter (lower limit=0.18, upper limit=0.28), and total spinal cord (lower limit=0.00, upper limit=0.29).

The automatic segmentation tool (radius of 2.5) was initiated on the thresholded image at the rostral and caudal most slices. The segmentation parameters were: competition force =1.000, smoothing force = 0.2, α=1.000, β=0.100, speed = 1.00. All parameters of contrast and thresholding were identical for each spinal cord. The mask was exported as a screenshot series of the transverse views and images every 100μm were quantified in FIJI 9 (U.S. National Institutes of Health). For spared white matter area, a custom macro was generated which: converted the series to a stack, isolated the region corresponding to the segmented ROI, and measured the total area for each slice. Volume for each parameter was calculated as the thickness of the voxel multiplied by the cross-sectional area. The cross-sectional area of the tissue occurred in a region 4.1mm in length and encompassing the epicentre.

### Immunohistochemistry for VGlut1

Following *ex vivo* MRI, spinal cords were washed in PBS, cryoprotected in 30% sucrose, embedded and frozen in OCT compound and cut in transverse 15μm sections. Immunohistochemistry for vGlut1 puncta was performed. Briefly, sections were immunostained using a rabbit anti-vGlut1 antibody (1:1000, Synaptic Systems) and Alexa Fluor 488 secondary antibody (1:1000, Life Technologies). 5 sections through the T7-T9 segments were imaged (LSM 710, Zeiss). Successfully traced α-motor neurons (Ctb+/ NeuN+) in the ventral horns were imaged at 400x magnification. Animals that had less than five successfully traced motor neurons were excluded from subsequent analysis.

#### Analysis

Analysis was performed in imageJ by measuring raw integrated density within a 2.5μm band surrounding the motor neuron, and manually counting distinct vGlut1+ puncta abutting the motor neuron.

### Statistical analysis

Prior to all experiments, power analysis (G*Power) was conducted to ensure sample sizes used were sufficient to yield reliable data based on previous experimental data to determine the expected effect size, type 1 error threshold (α) ≤ 0.05 and power (1-β) ≥ 0.90). Statistical analysis was performed using either Prism 9 (GraphPad) or SPSS (IBM). Locomotion behavioural assessments were analysed using a linear model with a covariate structure best suited to the data being analysed (Duricki et al., 2016b). All data sets were found to be normally distributed by viewing frequency histograms and satisfying Kolmogorov Smirnov and Shapiro-Wilk tests. Individual tests associated with each statistical analysis (and whether baseline values were included as a covariate) are explicitly mentioned in the figure captions. Error bars in graphs are mean ± Standard error of mean, and averages mentioned in text are mean ± standard deviation. Divergences were considered significant if P<0.05.

One animal from the NT3 group was excluded from the study, prior to the blinding being broken, based on consistently high scores within behavioural tests e.g. performing consistent weight-supported plantar steps within 7 days post injury and showing no discernible locomotive deficit at 21 days post injury.

## Results

### NT3 treatment recovered skilled motor function and coordination following injury

A contusion injury of 250kdyn at the T9 spinal level produced a severe SCI in 23 adult female Wistar rats. Animals were randomised into either an NT3 group (n=11, receiving a total of 2.57 x10^12^ GC of AAV-NT3 in 220μl PBS) or a control group (n= 12, receiving 220μl PBS). Uninjured rats served as the sham control (sham group, n=4) and received neither the displacement nor the muscle injections. No differences in the force or displacement of the spinal cord were observed between the control and treatment groups (Unpaired T test, Force of injury p=0.63, t=0.4899, df=21; Tissue displacement p=0.86, t=0.1757, df=21). Following injury, we performed a series of behavioural assessments (BBB, horizontal ladder, and swimming) to determine whether NT3 treatment improved locomotor function as compared to controls (experimental timeline, Figure 1). While the contusion injury significantly impacted the animals’ bilateral hindlimb locomotion across the open field 1 week following trauma (BBB; Figure 2A; Supplementary Figure 1A; effect of group, linear model, F(2,24)=106, p<0.0001, post hoc LSD comparison, NT3 vs Naïve p<0.0001, PBS vs Naïve p<0.0001), we found that AAV-NT3 treatment did not induce improved gross locomotive function compared to injured controls (Figure 2A; post hoc NT3 vs PBS p=0.92) nor was there an interaction (effect of time x group, linear model, F(18,213)=1.2, p=0.24). Indeed, both groups showed a plateau in recovery at three weeks post injury (3 wpi vs 10 wpi p=0.80, subsequent weeks vs 10wpi p>0.09). Assessment of total BBB subscores, showed a consistent but non-significant trend for the NT3 group to exhibit higher average functional capacity for both limbs compared to controls (Supplementary Figure 1B-C). NT3 treated animals tended to do modestly better than controls at all timepoints at BBB constituent components such as toe clearance (effect of group, linear model, F(2,24.3)=135, p<0.0001, post hoc LSD comparison, NT3 vs Naïve p<0.0001, PBS vs Naïve p<0.0001, NT3 vs PBS p=0.19) and paw position (Supplementary Figure 1D-F; effect of group, linear model, F(2,24.7)=63.2, p<0.0001, post hoc LSD; NT3 vs Naïve p<0.0001, PBS vs Naïve p<0.0001, N3 vs PBS p=0.13). Further, NT3 treatment improved trunk stability at later time points of nine and ten weeks post injury (Figure 2B; post hoc LSD NT3 vs PBS p=0.090 but with an interaction between group and time post injury, linear model, F (18,213)=2.78, p=0.0002, post hoc LSD NT3 vs PBS at 9 wpi p=0.024, at 10 wpi p=0.041). These data suggested that NT3 treatment may affect aspects of skilled hindlimb function rather than gross functionality.

**Figure 2:**
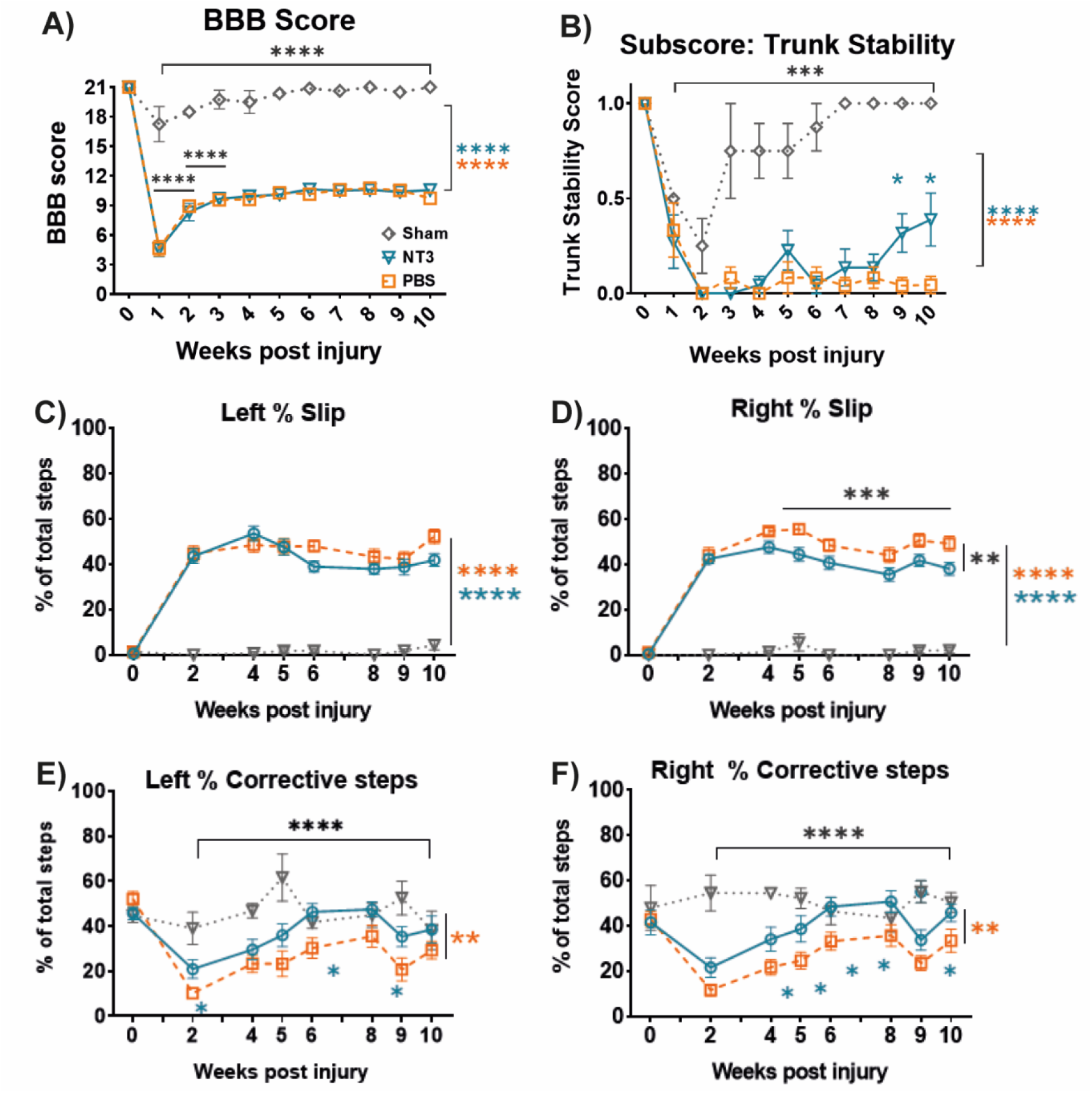
NT3 treatment can restore specific, skilled hindlimb functions. A) T9 contusion reduced the BBB score of all animals. NT3 treatment failed to cause recovery in total locomotive function compared to controls. Sham injured animals showed no sustained deficit in hindlimb function. B) Analysis of BBB sub-scores showed that trunk stability was impaired after injury compared to sham (effect of group, linear model, F(2,24.1)=75.2, p<0.0001, post hoc LSD; NT3 vs Naïve p<0.0001, PBS vs Naïve p<0.0001). NT3 treatment showed increases in trunk stability at 9 and 10 weeks post injury compared to PBS controls. C-D) NT3 treated animals showed a similar proportion of slips on the horizonal ladder with the C) left hindlimb and a decrease from 4 weeks post injury with the D) right hindlimb. Similarly, E-F) both hindlimbs of NT3 treated animals showed an increase in skilled corrective stepping following misplacement which did not disrupt locomotion compared to control groups. This functionality was comparable to sham animals. For panels C-F) Baseline recordings were not used as a covariate but were excluded for tests of effects and interactions. For NT3 n=11 for PBS n=11, for sham n=4. In all panels: NT3 = blue triangle, controls = orange square, sham = black diamond, and wpi = week post injury. * = P<0.05, ** = P<0.01, *** = P<0.001, and **** = P<0.0001. If no post-hoc result is shown, comparison was not-significant. Values represent mean±SEM.

In order to assess the degree of skilled hindlimb functional restoration following NT3 treatment animals were assessed on the horizontal ladder from two weeks post injury (Figure 2C-F). Contusion caused a sustained drop in the proportion of error free steps by both hindlimbs (Supplementary Figure 2A-B; effect of group, linear model, Left = F(2,23)=910, P<0.0001, Right = F(2,24)=693.6, P<0.0001) and increased the number of missed steps (Supplementary Figure 2C-D; effect of group, linear model, Left = F(2,24)=9.95, P=0.0007, post hoc LSD for NT3 vs Sham P=0.0053, PBS vs Sham P=0.0002, NT3 vs PBS p=0.079, Right = effect of group, linear model, F(2,24)=5.68, P=0.0095, post hoc LSD for NT3 vs Sham P=0.017, PBS vs Sham P=0.0025, NT3 vs PBS p=0.30). NT3 treatment did not increase the number of error free steps (NT3 vs PBS p=0.40) nor the number of missed steps (NT3 vs PBS at all timepoints p>0.15). However, NT3 treatment did improve the accuracy of hindlimb stepping on the right although this was not significant on the left (Figure 2C-D; effect of group, linear model, left = F(2,24)=120.6, P<0.0001, post hoc LSD for NT3 vs Sham P<0.0001, PBS vs Sham P<0.0001, NT3 vs PBS p=0.12, right = F (2, 24) = 111.5, P<0.0001, post hoc LSD for NT3 vs Sham P<0.0001, PBS vs Sham P<0.0001, NT3 vs PBS p=0.0027). Further, NT3 treated animals showed more frequent corrections following a misplaced step than controls, recovering functionality in this skilled task to the level of Sham injured animals from 6 weeks post injury (Figure 2E-F; effect of group, Linear model, Left = F(2,24)=6.75, P=0.0047, post hoc LSD for NT3 vs Sham p=0.13, PBS vs Sham P=0.0024 and NT3 vs PBS p=0.019, Right = F(2,24)=7.6, P=0.0026, post hoc Fisher’s LSD for NT3 vs Sham p=0.089, PBS vs Sham P=0.0012 and NT3 vs PBS p=0.016). These data suggested that NT3 treatment could enhance specific aspects of hindlimb function to aid skilled or coordinated functionality, potentially with tasks that involve proprioceptive feedback as would occur with corrective stepping or avoidance of slipping.

### NT3 treatment restores coordinated limb movement during swimming

To further assess the ability of NT3 treatment to facilitate skilled or coordinated motor tasks animals were assessed during an 80cm swim in a customised tank at a constant 20-23 °C to avoid temperature-evoked spasms (Figure 3A) (Gonzenbach et al., 2010). At baseline, animals consistently used the hindlimbs for propulsion in swimming and forepaws were kept in close contact to the body near the jaw (Figure 3Bi). Contused animals were able to swim but showed abnormal use of the limbs. Forelimbs and hindlimbs were used for propulsion with the tail submerged (Figure 3Bii). Contusion affected trunk stability, with the pelvic girdle rotated around the rostral caudal axis. These abnormalities persisted even when using the tank walls for stability (Figure 3Biii-iv).

**Figure 3:**
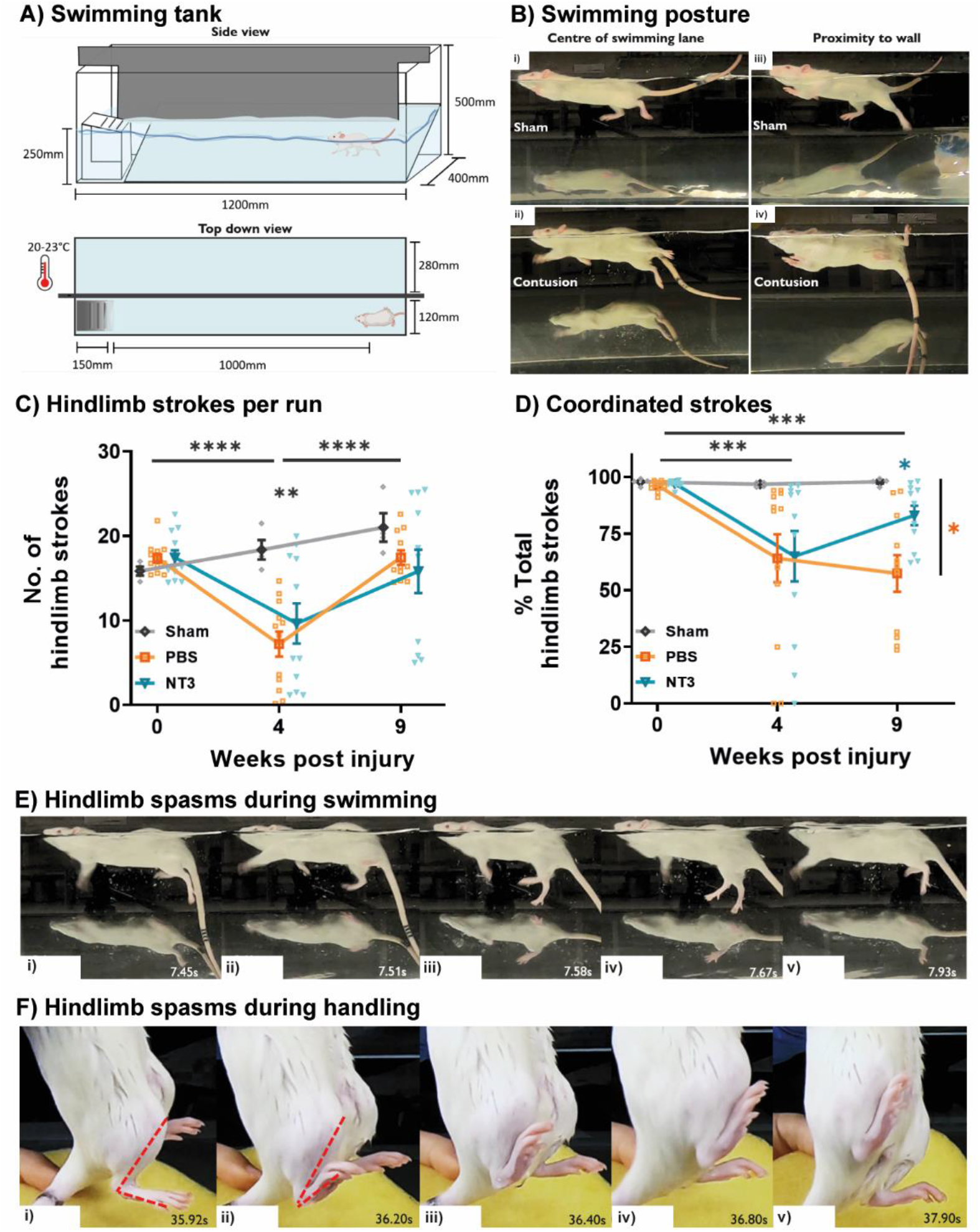
NT3 treatment restores deficits in hindlimb coordination during swimming. A) Swimming tank schematic. A 45°mirror provided simultaneous lateral and dorsal views. Water temperature = 20-23°C. B) Representative images showing swimming posture between i+iii) sham and ii+iv) contused animals (3 wpi) either i+ii) in the swimming lane or iii+iv) close to the tank walls (Supplementary Videos 1 & 2). C) The total sum of hind limb strokes, performed by both left and right hindlimbs. D) The proportion of alternating left:right hindlimb strokes following treatment. E) Representative images showing a 0.5 sec bilateral hindlimb spasms during open swimming where i) cyclic hindlimb strokes are ii) abruptly stopped, iii-iv) limbs extended, and then v) normal function resumes (Supplementary video 3). F) Representative images showing the progression of a 3 sec unilateral dorsiflexion spasm of the right hindpaw following swimming. Dotted red line i-ii) shows the closure of the angle made between the dorsal surface of the hindpaw and the shin. Adduction of the hip joint was also present, shown by iv) the inward rotation of the right hindlimb.

The effect of injury on swimming gait are reflected in the quantification of animals’ swimming activity. There was an ~50% reduction in the number of hindlimb strokes that animals performed during swimming 4 weeks after injury (Figure 3C; baseline: 16.6±2.2, NT3: 9.6±7.8, PBS: 7.2±5.1, and Sham: 18.3±2.3 strokes) which predominantly recovered by 9 weeks post injury (NT3: 15.9±8.8, PBS:17.4±2.8, Sham: 21.0±3.3 strokes; effect of time, linear model including all groups, F(1.52,35.8)=13.32, p=0.0002, post hoc LSD, baseline vs 4wpi p<0.0001, 4wpi vs 9wpi p<0.0001, baseline vs 9wpi p=0.92). However, at no stage was there a difference between treatment groups (effect of group, linear effects model, F(2,24)=2.13, p=0.15). Contusion injury caused a marked reduction in hindlimb coordination, with a lower proportion of strokes following left-right rhythmicity (Figure 3D; effect of time, linear model, F(1.52,35.75)=7.15, p=0.005, post hoc LSD, baseline vs 4wpi p=0.0003, baseline vs 9wpi p<0.0001, 4wpi vs 9wpi p=0.36)). Overtime, NT3 treatment caused an increase in the proportion of coordinated strokes such that activity was similar to that of sham animals at 9 weeks, whilst PBS treated animals had no improvement (effect of group x time, linear model F(4,47)=2.57, p=0.48, post hoc LSD at week 9, NT3 vs PBS p =0.013; NT3 v sham p=0.272; and PBS v sham p=0.0041). These data similarly suggest that NT3 treatment could enhance specific aspects of hindlimb function aiding coordinated movement.

### Contusion did not cause hyperreflexia of the propriospinal circuity

Only one animal showed noticeable spasm, clonus or otherwise abnormal activation of the hindlimbs or trunk during swimming (during one single instance) (Figure 3E) and only five animals showed presumptive spasm activity once following swimming (Figure 3F). This was unexpected as this is activity typically associated with this injury (Ryu et al., 2021).

To investigate hyperreflexia, the H reflex of the animals was recorded at 11 weeks post injury by stimulation of the medial plantar nerve over a defined range of frequencies following all locomotor testing (Figure 4A-B) (Smith et al., 2018, Lee-Kubli and Calcutt, 2014, Chang et al., 2019, Kathe et al., 2016b). The amplitude of the H wave remained comparable in all three groups (NT3=103.1 ±5%, PBS 102.2 ±2.6%, sham 101.6 ±1.3%; Figure 4C-E). Animals were excluded from analysis if they received a higher dose of anaesthetic prior to recording or limited rate dependent depression (RDD) from 2000ms to 50ms (as occurred in one sham animal). This did not affect the conclusions drawn (Supplementary Figure 3A-F).

**Figure 4:**
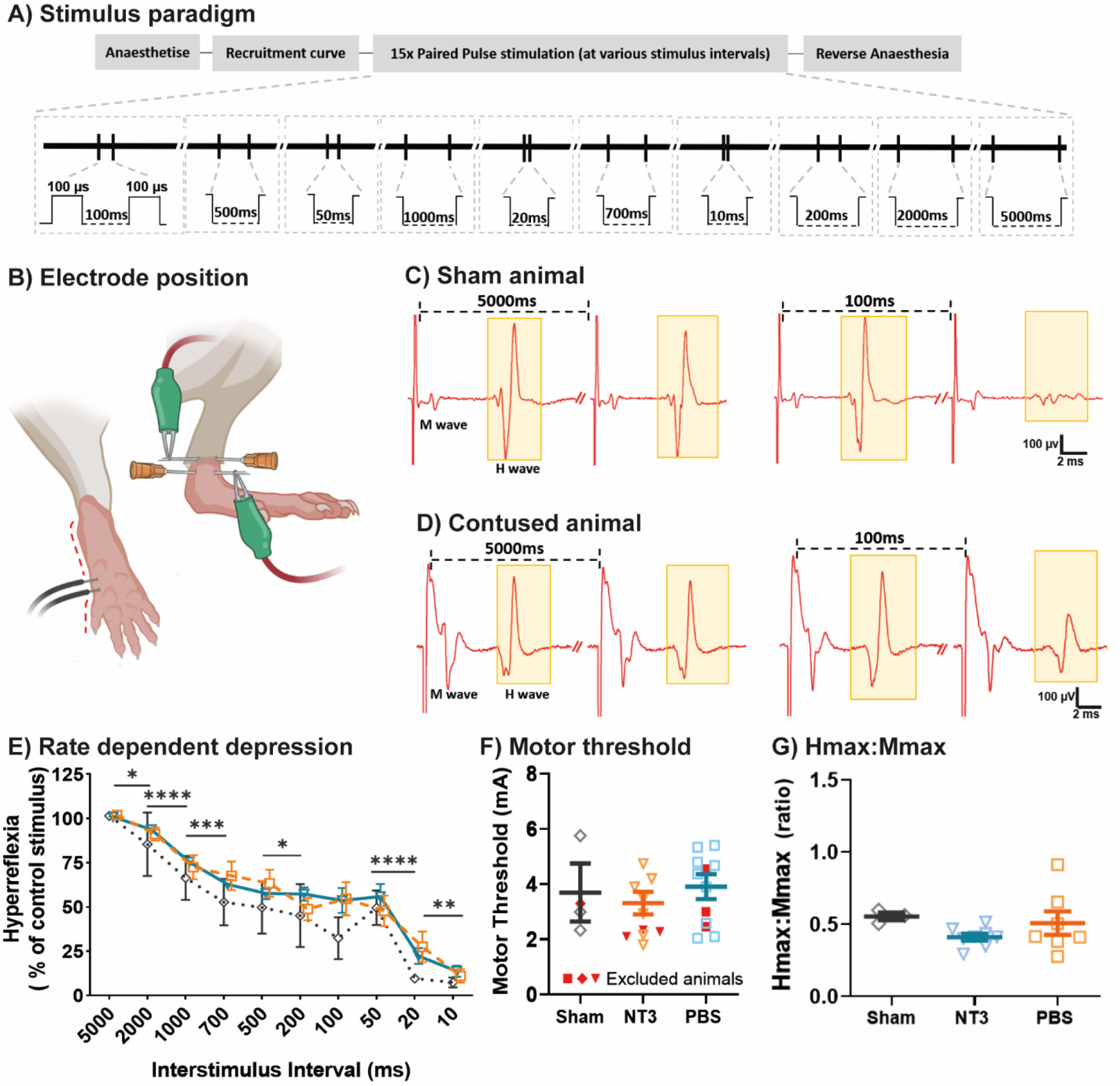
250kdyn thoracic contusion did not generate hyperreflexia. A) Stimulation protocol. B) Position of the recording and stimulation electrodes. C&D) Representative traces from C) sham and D) contused animals highlighting the M and H wave (yellow). E) Rate dependent depression was the same regardless of animal group. F) Motor threshold highlighting those from excluded animals. G) The Hmax:Mmax ratio was comparable between groups (One way ANOVA, F(2,15)=1.43, p=0.23).

RDD was comparable between injured and sham groups (Figure 4E; effect of group, linear model, F(2,15)=0.83, p=0.45). RDD increased with shorter interstimulus interval (effect of interstimulus interval, linear model, F(1.9,27.6)=64.1, P<0.0001, 5000ms vs 2000ms p<0.0001, 2000ms vs 1000ms p=<0.0001, 1000ms vs700ms p=0.0007, 700ms vs 500ms p= 0.095, 500ms vs 200ms p=0.021, 200ms vs 100ms p=0.73, 100ms vs 50ms p=0.81, 50ms vs 20ms p<0.0001, 20ms vs 10ms p=0.0046). Sham animals had a consistent trend towards smaller H waves amplitudes at each interstimulus interval. The motor threshold was not altered following injury or treatment (Figure 4F; one way ANOVA, F(2,17)=0.42, p=0.65). The maximum H wave amplitude, Hmax, and maximum M wave amplitude, Mmax, remained comparable amongst all groups (Supplementary Figure 3G-H). The ratio between the Hmax and Mmax values, which can be used as an indicator for the excitability of the reflex, was unaltered after injury (Figure 4G). Overall, these results indicate that a lumbar H reflex was not detectably altered by severe thoracic contusion compared to shams.

### Intramuscular injection of AAV1-CMV-NT3 elevated NT3 serum levels

Serum was collected from all animals 11 weeks following injury and processed using ELISA to quantify levels of circulatory NT3 protein. Results were expressed as a proportion of the total protein content in each sample. Injection of AAV-NT3 into the gastrocnemius, tibialis anterior and rectus abdominus muscles resulted in elevation of NT3 levels in serum (Figure 5A; NT3 = 9.05±0.61 pg/mg, PBS = 0.12±0.004 pg/mg, Sham = 0.13±0.003 pg/mg; effect of group, one way ANOVA, F(2,23)=143.2, p<0.0001, post hoc LSD NT3 vs PBS p<0.0001, NT3 vs Naïve p<0.0001). This demonstrated that the AAV was producing NT3.

**Figure 5:**
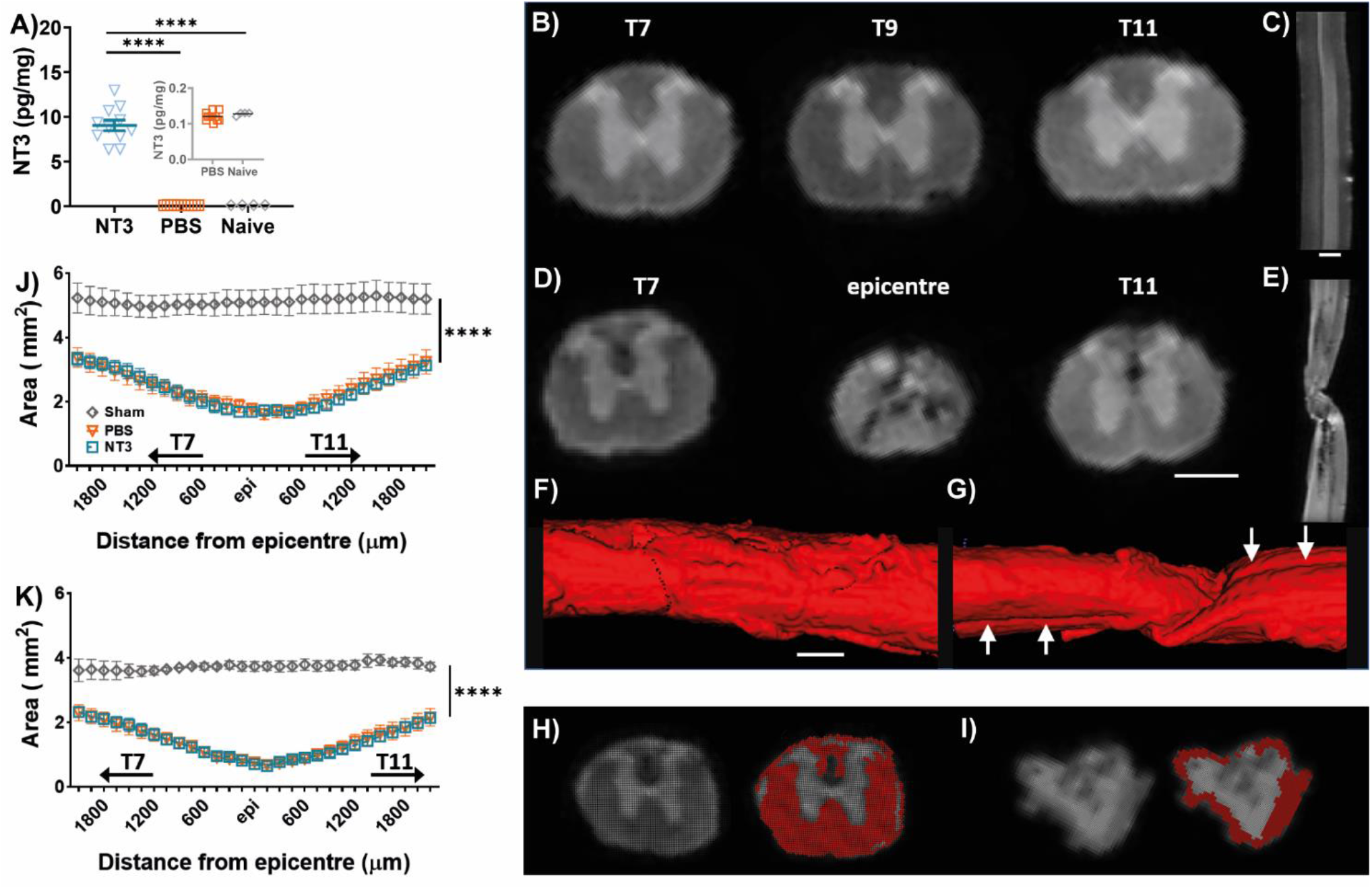
Intramuscular injection of AAV-NT3 increased circulating NT3 levels but did not alter lesion volumetrics. A) Intramuscular injection of AAV-NT3 resulted in an elevation of NT3 protein in serum in all treated animals. SCI followed by intramuscular injections of PBS did not alter the level of NT3 in serum, shown in insert (post hoc LSD, PBS vs Naïve p=0.99). B) Transverse sections through an intact spinal cord at T7, T9, and T11, showing the characteristic grey and white matter morphology. C) Sagittal view of the intact cord. D) Transverse sections of a chronically contused spinal cord at T7, T9, and T11 demonstrating hypointense imaging in the lesion epicentre. E) Sagittal section of the injured cord where the lesion epicentre apparent from the reduced width of the spinal cord and loss of grey/ white matter distinction. Hypointense signal was seen extending caudally from the epicentre. F&G) Sagittal, volumetric renderings of the F) intact and G) injured spinal cords (T7 through T11). Twisting of the epicentre was visualised by tracing the ventral fissure, indicated by arrow, as it courses caudally through the lesion. H&I) T2 Weighted MR images showing H) T7 and I) the epicentre with the corresponding automatically generated mask of presumptive spared white matter. J) Total cross-sectional area of the spinal cord, inclusive of spared tissue and cavity, throughout the lower thoracic spinal cord. K) Presumptive spared white matter through the thoracic spinal cord following treatment.

### NT3 treatment did not affect lesion volume

*Ex vivo* Magnetic Resonance Imaging (MRI) was performed following termination of the experiment 11 weeks after thoracic contusion to assess any differences in lesion volume between treatment and control groups. This generated high resolution T2 weighted images of the thoracic spinal cord (Figure 5B-I). At this resolution, there was a clear distinction the grey and white matter from T7 to T11 on both sham (Figure 5B-C&F) and contused cords (Figure 5D-E&G). The epicentre was determined as the spinal cord transverse section with the smallest cross-sectional area of presumptive spared white matter. Within the epicentre there was a mixture of fragmented hypointense signal, likely corresponding to fluid filled cavitation, interspersed with scar tissue and/or oedema. Further, sagittal sections showed that the injured spinal cords were deformed at the contused region, a likely artefact of imaging which arose from insertion of the cords into the immobilisation device prior to imaging (Figure 5E&G). These two factors make establishing cavity volumes difficult.

Total spinal cord transverse cross-sectional area and presumptive spared white matter were measured following thresholding for the respective tissue and automatic detection in ITK-SNAP software. Presumptive spared white matter was defined as regions with a T2 weighted signal comparable to, and continuous with, the signal in the lateral funiculus in the rostral and caudal most spinal level imaged (Figure 5A&H-I). The contusion injury was shown to reduce the total transverse cross-sectional area extending at least 2mm rostral and caudal from the lesion epicentre (Figure 5B&J; effect of group, two way RM ANOVA, F(2,23)=29.41, p<0.0001; NT3 vs sham p<0.0001, PBS vs sham p<0.0001). Overall total transverse cross-sectional area remained similar in both treatment groups (post hoc LSD NT3 vs PBS p=0.86). On average, the total transverse cross-sectional area at the epicentre was reduced by 68% in the NT3 group compared to the equivalent spinal cord level in sham (NT3=1.68 ±0.5 mm^2^, PBS=1.61 ±0.5 mm^2^ and sham=5.10±0.5 mm^2^; effect of group x level, two way RM ANOVA, F(56, 637)=2.65, p<0.0001, rostral most region; NT3 vs naïve p<0.0001, PBS vs naïve p<0.0001, at epicentre; NT3 vs naïve p<0.0001, PBS vs naïve p<0.0001 and caudal most region NT3 vs naïve p<0.0001, PBS vs naïve p<0.0001). Further, the contusion injury reduced the amount of presumptive spared white matter in the caudal thoracic cord (Figure 5C; effect of group, two way RM ANOVA, F(2,22)=47.28, P<0.0001, post hoc LSD NT3 vs sham p<0.0001, PBS vs Naïve p<0.0001). On average presumptive spared white matter was reduced equally in the epicentres of both PBS and NT3 treated groups, where it was reduced by 36% compared to sham (NT3=0.64±0.17 mm^2^, PBS=0.67±0.30 mm^2^ and sham=3.7±0.2 mm^2^; post hoc LSD NT3 vs PBS p=0.94). Presumptive spared matter remained reduced 2mm rostral and caudal from the epicentre (effect of group x level, two way RM ANOVA, F(56,612)=2.49, p<0.0001, post hoc LSD rostral most region NT3 vs naïve p<0.0001, PBS vs naïve p<0.0001; at epicentre, NT3 vs naïve p<0.0001, PBS vs naïve p<0.0001; and caudal most region NT3 vs naïve p=0.0002, PBS vs naïve p<0.0001). These data would suggest that NT3 treatment did not facilitate in reducing lesion volumes (consistent with the 48 hour delay in treatment utilising a viral vector).

### NT3 normalised vGlut1+ excitatory input onto motor neurons

To determine treatment induced proprioceptive sprouting, the Ia afferent fibres making direct glutamatergic (vGlut1 +) connections with α-motor neurons were assessed through immunohistochemistry. The motor neurons innervating the lateral head of the *gastrocnemius* were retrogradely traced with Cholera Toxin B (CtB, Figures 1A&6A), injected intramuscularly and targeting the muscle spindles and motor end plates (Yin et al., 2019). The L4-L6 spinal cord region, encompassing the *gastrocnemius* motor pool (Mohan et al., 2014), was stained for vGlut1 (Ia afferent derived motor neuron input) and for CTb (motor neurons innervating the lateral head of the *gastrocnemius*) (Figure 6B). In order to exclude afferent innervation onto homonymous γ-motor neurons, the mature neuronal marker NeuN was used to distinguish between α- and γ-motor neurons (Friese et al., 2009).

**Figure 6:**
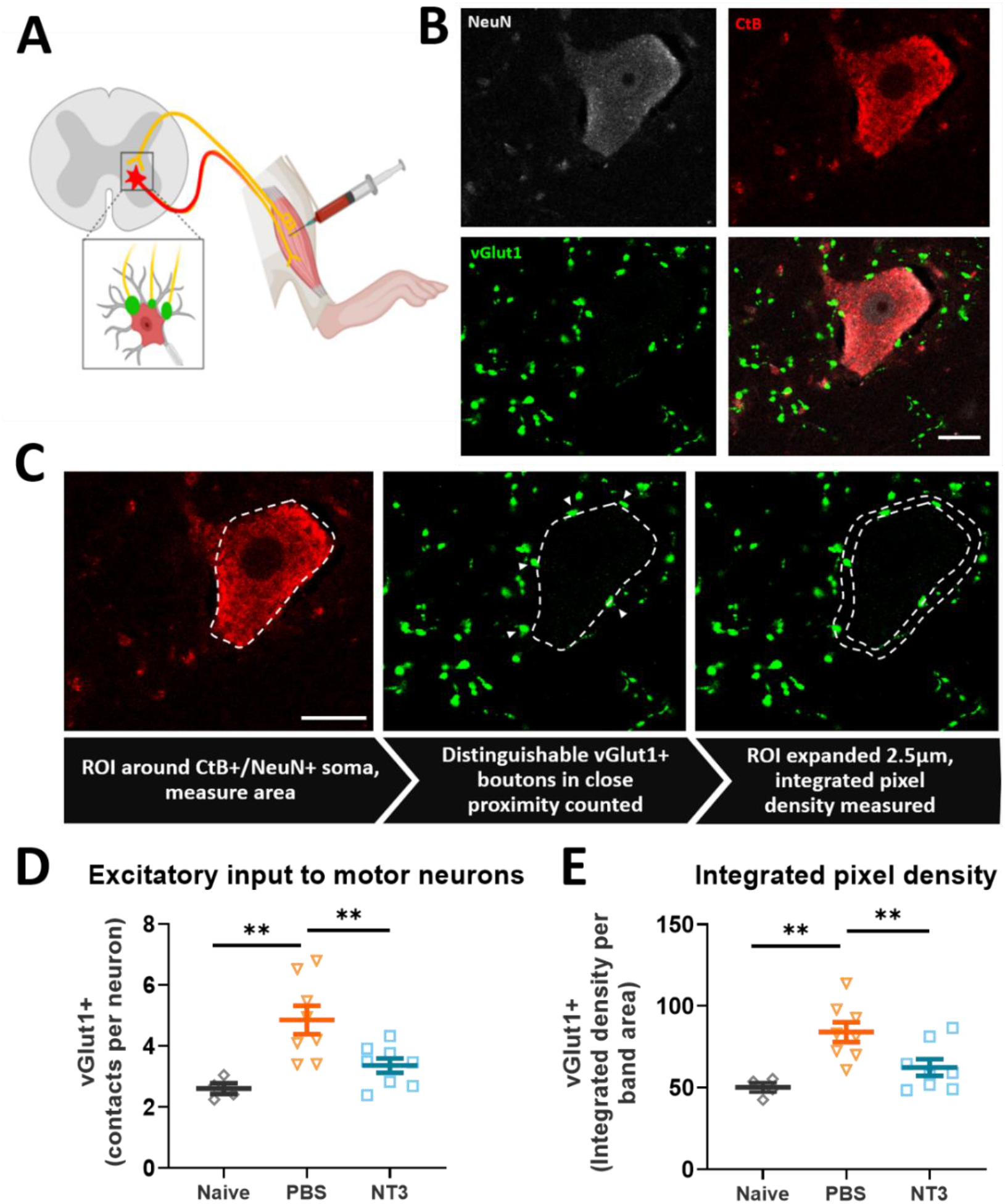
Injury induced afferent derived excitatory input was normalised by NT3 treatment. A) Motor neurons innervating the lateral head of the left *gastrocnemius* were retrogradely traced with intramuscular CTb injection to target the motor end plates. Ia afferent fibres innervating the lateral *gastrocnemius* make direct vGlut1 + contacts with these homonymous motor neurons. B) Numerous vGlut1 puncta were present on both the soma compartment and proximal dendrites of traced α-motor neurons (CTb+/ NeuN+). C) Overview of the methods used to quantify vGlut1 + puncta on motor neuron. Arrow heads mark distinct vGlut1 puncta which were abutting the soma membrane and counted. D) T8 contusion increased individual vGlut1+ puncta abutting CTb+/NeuN+ soma, which was normalised back to levels comparable with uninjured animals by NT3 treatment. E) NT3 was able to normalise the increase in integrated density of vGlut1+ signal within a 2.5μm band around the soma after thoracic contusion. For all panels, scale bar = 20μm. Animals per group: Naïve =4, NT3 & PBS=8. Total number of motor neurons analysed: Naïve=58, NT3= 111 and PBS=117, Number of Puncta: Naïve=153, NT3=338 and PBS=556.

To account for the varying sizes of vGlut1+ immunoreactive puncta seen on the motor neuron perimeter, two quantitative approaches were taken: 1) as individual counts and 2) as integrated pixel intensity, which incorporated the varying size and brightness of the vGlut1 signal. Using the former quantification method, a near doubling of vGlut1+ puncta close to the motor neuron soma was observed following contusion injury (average puncta in Naïve=2.6±0.17, PBS=4.9±0.45 puncta; Figure 6D). However, NT3 treatment was able to reduce this increased excitatory input back to a level comparable with naïve animals (NT3 group: 3.4±0.23 puncta; one way ANOVA, F(2,17)=8.75, P=0.0024, post hoc LSD; NT3 vs PBS p=0.0063, NT3 vs Naïve p=0.21 PBS vs Naïve p=0.0014). A similar result was detected taking into account integrated pixel intensity (Figure 6E; one way ANOVA, F(2,17)=8.47, P=0.0028, post hoc LSD; NT3 vs PBS p=0.0083, NT3 vs Naïve p=0.19 PBS vs Naïve p=0.0014). These results indicate that intramuscular NT3 treatment normalised the number of vGlut1 + puncta on hindlimb motor neurons after contusion injury.

## Discussion

The aim of this study was to determine whether a clinically relevant delayed delivery of NT3 to the muscles of the ankle, *tibialis anterior* and *gastrocnemius*, and lower portion of the *rectus abdominus* following a T8 contusion would modulate the lumbar neural circuitry facilitating functional recovery. We demonstrated that treated animals had a 75 fold increase in circulating NT3 levels at 11 weeks post injury. This did not affect the size of the injury produced (consistent with delayed treatment application). Further, these animals did not demonstrate gross improvement in behavioural function but showed through numerous motor tasks that the treatment strategy enhanced skilled hindlimb placement in walking, coordination of these limbs when swimming, and trunk stability, likely involving proprioceptive feedback. We suggest that this increase in function is due to the normalisation of vGlut1+ boutons on presumptive proprioceptive afferents innervating these muscles. Interestingly, the injury did not result in measurable signs of spasticity, such as hindlimb spasms and hyperreflexia, suggesting theses typical SCI outcomes are not reliably induced through severe thoracic contusion. This work is the first demonstration that a clinically relevant 48-hour delay in AAV1-NT3 treatment application can help restore fine functional motor control, supporting its use as a therapeutic strategy for SCI.

We used an AAV1-mediated gene therapy to deliver NT3 via injection into peripheral muscle. This has the benefit of being an effective and safe delivery system (Fortun et al., 2009, Petruska et al., 2010) *(e.g.,* as used in Glybera for another transgene) (Sydney-Smith et al., 2021b, Kakanos and Moon, 2019, Wang et al., 2018, Zhou et al., 2003, Chen et al., 2006). Neurotrophin-3 is safe and well-tolerated in human adults when given peripherally at high doses (Sahenk et al., 2005, Parkman et al., 2003, Coulie et al., 2000). Here we confirmed that the AAV serotype 1 could produce high quantities of circulating NT3 long after initial injection following delivery; we and others have shown that NT3 is delivered to DRG and to motor neurons by retrograde transport of the protein from muscle after intramuscular or intraneural injection of AAV1-CMV-NT3 (Curtis et al., 1998, DiStefano et al., 1992, Kathe et al., 2016b, Han et al., 2019, Wang et al., 2018).

### NT3 treatment promoted recovery in skilled control of the hindlimb

Our results showed a beneficial effect of NT3 treatment on trunk stability, swimming coordination, and skilled hindlimb placement in walking. The contusion produced a severe effect on hindlimb motor function which was not recovered up to 11 weeks post injury as demonstrated through the BBB where, even following endogenous recovery, the animals were unable to regain consistent plantar stepping and interlimb coordination. These data are consistent with previous studies following moderate thoracic contusion from a weight drop mechanism (Bose et al., 2012, Basso et al., 1995). The locomotor impairments we report were, unexpectedly, not as severe as studies which also utilised a 250 kdyn contusion at either T7 (Park et al., 2016) or T8 (Chang et al., 2019, Ryu et al., 2021) spinal level. In these studies, injured animals had limited hindlimb movement one week after injury and plateaued in recovery from week four showing the ability to perform dorsal stepping and plantar placement in stance but not weight supportive plantar stepping during movement (e.g., at BBB=9 in Chang et al 2019 versus BBB=10 in ours). These subtle differences between our study and theirs in functional performance following injury may suggest the contusion performed here did not induce as severe a neurological deficit as expected despite similar injury parameters (e.g., actual force delivered).

Data regarding toe clearance and trunk stability was of particular significance to the study as NT3 was applied to the main ankle flexor and extensor muscles and the rectus abdominus muscles. Whilst there was no difference between groups in toe clearance, NT3 treated animals did consistently score averages above PBS treated rats. Notably, here we show an effect of NT3 treatment on trunk stability during open field walking from seven weeks post injury. Trunk and pelvic support involves numerous muscles, including; the *rectus abdominus, obliques* and muscles of the vertebral column such as *erector spinae* (Musienko et al., 2014). Here only the caudal half of the *rectus abdominus* was injected with AAV-NT3 (although NT3 circulating in the blood might have affected non-injected muscles). Similar effects upon trunk stability have been achieved in previous thoracic contusion experimental studies 8 weeks post injury following regular training up and down stairs (Singh et al., 2011). These data may suggest that NT3 treatment can mimic some of the effects of regular training in specific circumstances. Of course, trunk stability was not improved during swimming within our study, however, this could be due to the lack of weight bearing/ the effects of buoyancy during swimming. However, this could be due to the lack of weight bearing during swimming. This difference in experimental condition may exaggerate any instability that is masked during over ground locomotion. Further experimentation where all muscles involved in trunk stability are injected with NT3 should be considered (*obliques*, *transversus abdominis and/*or *erector spinae*).

The effect of NT3 improving precision hindlimb stepping on the horizontal ladder is consistent with other work (Duricki et al., 2016a, Kathe et al., 2016b, Fortun et al., 2009). Administration of a viral vector encoding NT3 into the sciatic nerve partially recovered error free hindlimb stepping after a thoracic contusion at T9 (Han et al., 2019). They showed the contusion generated atrophy of dendrites on motor neurons caudal to the injury and associated with loss of synaptic connections. Elevation of NT3 within lumbar motor neurons (retrogradely transported from the nerve) encouraged regrowth of these previously atrophied motor neuron dendrites. This was accompanied by elevated synaptic like contacts between long descending propriospinal neuron terminals and motor neuron soma. As retrograde transport of NT3 occurs from peripheral tissues to motor neurons (DiStefano et al., 1992), it is possible the recovery we see in hindlimb performance is due to a similar mechanism: NT3 may induce recovery after secretion from motor neurons by inducing neuroplasticity of descending axons.

Intramuscular injection of AAV-NT3 to forelimb flexors in female Lister Hooded rats after bilateral pyramidotomy injury reduced the number of stepping errors during locomotion (Kathe et al., 2016a). NT3 protein accumulated in cervical DRGs most likely a result of transport of NT3 within Ia afferents. RNA-seq identified a number of transcriptional changes occurring within the DRG in response to NT3 treatment (Kathe and Moon, 2018). These include upregulation of *Sema4c*, which belongs to a family of axon guidance molecules that work through attraction or repulsion mechanisms, and cytoskeletal remodelling associated genes like *Gas7*. Such transcriptional changes may stabilise afferent contacts on motor neurons and strengthen proprioceptive circuitry involved in the stretch reflex, reflected by the reduction of vGlut1+ afferent derived input on homonymous motor neurons (Kathe et al., 2016b). NT3 treatment reduced afferent input onto *gastrocnemius* α-motor neurons in our study, suggesting recovery of precision ladder stepping may require, in part, normalisation of vGlut1+ input. Of particular note is the shift in the type of stepping error in NT3 treated animals in our study; from slips that hinder locomotion to corrective steps. This may be explained by a more efficient stretch reflex response to a poorly positioned paw or beginning of a slip, enabling repositioning of the paw (before transfer of weight support and slippage). As slipping would involve activation of Ia afferents innervating flexors and extensors of the ankle joints, it may be that this proprioceptive circuitry was abnormal after injury; however, note that in our study the H-reflex was recorded from the hindpaw muscle and not the ankle muscles).

NT3 treatment caused an increase in alternating hindlimb coordination during swimming. Following contusion, we report animals exhibited abnormalities of the swimming gait including; tail deflection below the water, trunk instability and frequent forelimb use (which did not occur in any of the baseline recordings) (Smith et al., 2006, Xu et al., 2015) which appeared to decrease following NT3 treatment. However, while the T8 contusion injury used would not have damaged the hindlimb central pattern generator circuitry located within the lumbar spinal cord (Langlet et al., 2005), based on the location of reduced presumptive spared white matter located throughout the lesion epicentre, the injury likely damaged some long descending propriospinal neurons. These interneurons relay afferent input allowing coordinated movement between fore and hind limbs (Brockett et al., 2013, Reed et al., 2006) and alternation of hindlimbs at locomotion speeds comparable to that during swimming via commissural projecting interneurons (Ruder et al., 2016). This is likely to occur through the excitatory and inhibitory V0 population of interneurons (Talpalar et al., 2013). A possible explanation for recovery of coordination during swimming with NT3 treatment may relate to the effects of retrogradely transported NT3 has on lumbar motor neuron dendritic regrowth following injury and the formation of descending detour pathways (Han et al., 2019). More extensive dendritic arbors after treatment would enable a greater amount of propriospinal input to be received, possibly from the contralaterally projecting propriospinal interneurons directly or the V0 population.

### T8 contusion model did not reliably induce spasms or hypereflexia

Our study reveals that hindlimb spasms are not reliably generated by a 250kdyn contusion at the T8 vertebral level in Wistar rats, contrasting with previous reports using the same injury model (Ryu et al., 2017, Gonzenbach et al., 2010, Chang et al., 2019). Reasons for this could be due to the strains used (Mills et al., 2001), or slight alterations in equipment or methodologies employed. Our injuries were not as functionally extensive as those previously reported with this injury, perhaps suggesting the CST was not as comprehensively injured and perhaps accounting for this effect (Gonzenbach et al., 2010). Alternatively, spared reticulospinal function is positively correlated with the presence of hindlimb spasticity in SCI patients (Sangari and Perez, 2019), suggesting we may have damaged this pathway too excessively. Indeed, the lack of hyperreflexia in our animals (Boulenguez et al., 2010, Bandaru et al., 2015) in addition to the lack of visual spasms would suggest that the T8 contusion used does not reliably yield stereotypical functional responses. It is possible that altering the methods employed may have yielded different data. For example, assessing modest and mild spasms through EMG recordings from affected muscles, or a train of stimuli to more fully depress the H reflex (Lee et al., 2014) rather than paired-pulses (Chang et al., 2019, Ryu et al., 2017). However, as the methods employed have been successful in a number of studies previously, this study highlights that production of visual spasms and hyperreflexia of the hindpaw is not as a reliable outcome of severe T8 contusive injuries as previously reported.

### Ex-vivo *imaging of the spinal cord showed an extensive injury was produced*

*Ex vivo* MRI revealed an extensive lesion, shown by reduced presumptive spared white matter several millimeters rostral and caudal from the epicentre caused by our injury. This metric was chosen as white matter sparing is a frequently measured indicator of lesion size and has been found to correlate with functional recovery. Quantification of experimental SCI by MRI also closely matches histologically assessed injury size (Ditor et al., 2008, Byrnes et al., 2010, Mihai et al., 2008). We report sparing of the lateral and ventral white matter including; the reticulospinal tract (Schucht et al., 2002), propriospinal neurons (Flynn, Conn et al. 2017; Sheikh, Keefe et al. 2018) and possibly the rubrospinal (Morris and Whishaw, 2016). These data would potentially explain the lack of spasms in our animals and the deficits shown in fine locomotor control and synchronised movements (Jordan et al., 2008). The corticospinal tract, which relays input for fine motor control, is primarily located in the dorsal columns which appear to have the least white matter sparing. Tracing experiments after a similar severity of contusion to the cervical spinal cord disrupts all but a few CST fibres in the dorsal lateral white matter (Anderson et al., 2009). Given the diameter of the lumbar cord is a third that of the cervical cord (Watson et al., 2009), it is unlikely more than a few of these CST fibres remain caudal to the injury.

## Conclusion

We show that targeted, delayed delivery of NT3 to many hindlimb and trunk muscles 48h following a severe contusive injury to the thoracic spinal cord aids fine locomotor control and synchronised movements in rats. We demonstrate that NT3 improved trunk stability, accuracy of stepping during skilled locomotive tasks, and alternation of the hindlimbs during swimming (likely involving proprioceptive feedback), but it had no effect on gross locomotion function in the open field. We suggest that this increase in function is due to the normalisation of vGlut1+ boutons on presumptive proprioceptive afferents innervating these muscles. Further, here we show that a T8 250kDyn contusion does not reliably result in measurable signs of spasticity, such as hindlimb spasms and hyperreflexia. This is the first demonstration that a clinically relevant 48-hour delay in AAV1-NT3 treatment application can help restore precise motor control in functionally compromised hindlimbs, supporting its use as a therapeutic strategy for SCI and continued clinical development.

## Additional information

### Data availability

Data is available from the corresponding author upon reasonable request.

### Author contributions

Animal work, data acquisition, tissue processing, immunohistochemistry, imaging, and data analysis were performed by J.D.S-S. and A.K., manuscript preparation was performed by P.M.W and L.D.F.M., and manuscript editing was performed by all the authors. The project was conceived and designed by P.M.W and L.D.F.M. in consultation with A.K. and J.D.S-S. All authors have approved the final version of the manuscript, agree to be accountable for the work, and qualify of authorship.

## Acknowledgements

The authors wish to thank Dr. Aline Spejo for her assistance with the ELISA experiments. We are grateful to the veterinary and technical staff at King’s College London for their proficiency and care of the animals. We acknowledge the Penn Vector Core in the Gene Therapy Program of the University of Pennsylvania for production of the AAVs for this project. We thank Dr Diana Cash and Dr Eugene Kim (King’s College London preclinical imaging core) for their help with acquisition of magnetic resonance images.

## References

Anderson, K. D., Sharp, K. G. & Steward, O. 2009. Bilateral cervical contusion spinal cord injury in rats. Exp Neurol, 220, 9–22.

Bandaru, S. P., Liu, S., Waxman, S. G. & Tan, A. M. 2015. Dendritic spine dysgenesis contributes to hyperreflexia after spinal cord injury. J Neurophysiol, 113, 1598–615.

Basso, D. M., Beattie, M. S. & Bresnahan, J. C. 1995. A sensitive and reliable locomotor rating scale for open field testing in rats. Journal of neurotrauma, 12, 1–21.

Bose, P. K., Hou, J., Parmer, R., Reier, P. J. & Thompson, F. J. 2012. Altered patterns of reflex excitability, balance, and locomotion following spinal cord injury and locomotor training. Front Physiol, 3, 258.

Boulenguez, P., Liabeuf, S., Bos, R., Bras, H., Jean-Xavier, C., Brocard, C., Stil, A., Darbon, P., Cattaert, D., Delpire, E., Marsala, M. & Vinay, L. 2010. Down-regulation of the potassium-chloride cotransporter KCC2 contributes to spasticity after spinal cord injury. Nature Medicine, 16, 302–307.

Boyce, V. S. & Mendell, L. M. 2014a. Neurotrophic factors in spinal cord injury. Handb Exp Pharmacol, 220, 443–60.

Boyce, V. S. & Mendell, L. M. 2014b. Neurotrophins and spinal circuit function. Front Neural Circuits, 8, 59.

Brockett, E. G., Seenan, P. G., Bannatyne, B. A. & Maxwell, D. J. 2013. Ascending and descending propriospinal pathways between lumbar and cervical segments in the rat: evidence for a substantial ascending excitatory pathway. Neuroscience, 240, 83–97.

Byrnes, K. R., Fricke, S. T. & Faden, A. I. 2010. Neuropathological differences between rats and mice after spinal cord injury. Journal of magnetic resonance imaging: JMRI, 32, 836–846.

Chang, Y.-X., Zhao, Y., Pan, S., Qi, Z.-P., Kong, W.-J., Pan, Y.-R., Li, H.-R. & Yang, X.-Y. 2019. Intramuscular Injection of Adenoassociated Virus Encoding Human Neurotrophic Factor 3 and Exercise Intervention Contribute to Reduce Spasms after Spinal Cord Injury. Neural Plasticity, 2019, 14.

Chaudhry, V., Giuliani, M., Petty, B. G., Lee, D., Seyedsadr, M., Hilt, D. & Cornblath, D. R. 2000. Tolerability of recombinant-methionyl human neurotrophin-3 (r-metHuNT3) in healthy subjects. Muscle & nerve, 23, 189–92.

Chen, Q., Zhou, L. & Shine, H. D. 2006. Expression of neurotrophin-3 promotes axonal plasticity in the acute but not chronic injured spinal cord. J Neurotrauma, 23, 1254–60.

Coulie, B., Szarka, L. A., Camilleri, M., Burton, D. D., Mckinzie, S., Stambler, N. & Cedarbaum, J. M. 2000. Recombinant human neurotrophic factors accelerate colonic transit and relieve constipation in humans. Gastroenterology, 119, 41–50.

Curtis, R., Tonra, J. R., Stark, J. L., Adryan, K. Park, J. S., Cliffer, K. D., Lindsay, R. M. & Distefano, P. S. 1998. Neuronal injury increases retrograde axonal transport of the neurotrophins to spinal sensory neurons and motor neurons via multiple receptor mechanisms. Mol Cell Neurosci, 12, 105–18.

Da Silva, S. & Wang, F. 2011. Retrograde neural circuit specification by target-derived neurotrophins and growth factors. Curr Opin Neurobiol, 21, 61–7.

Distefano, P. S., Friedman, B., Radziejewski, C., Alexander, C., Boland, P., Schick, C. Lindsay, R. M. & Wiegand, S. J. 1992. The neurotrophins BDNF, NT-3, and NGF display distinct patterns of retrograde axonal transport in peripheral and central neurons. Neuron, 8, 983–93.

Ditor, D. S., John, S., Cakiroglu, J., Kittmer, C., Foster, P. J. & Weaver, L. C. 2008. Magnetic resonance imaging versus histological assessment for estimation of lesion volume after experimental spinal cord injury. Laboratory investigation. J Neurosurg Spine, 9, 301–6.

Divanoglou, A. & Levi, R. 2009. Incidence of traumatic spinal cord injury in Thessaloniki, Greece and Stockholm, Sweden: a prospective population-based study. Spinal Cord, 47, 796–801.

Duricki, D. A., Hutson, T. H., Kathe, C., Soleman, S., Gonzalez-Carter, D., Petruska J. C., Shine, H. D., Chen, Q., Wood, T. C., Bernanos, M., Cash, D., Williams, S. C., Gage, F. H. & Moon, L. D. 2016a. Delayed intramuscular human neurotrophin-3 improves recovery in adult and elderly rats after stroke. Brain, 139, 259–75.

Duricki, D. A., Soleman, S. & Moon, L. D. F. 2016b. Analysis of longitudinal data from animals with missing values using SPSS. Nature Protocols, 11, 1112–1129.

Edgerton, V. R., Courtine, G., Gerasimenko, Y. P., Lavrov, I., Ichiyama, R. Fong, A. J., Cai, L. L., Otoshi, C. K., Tillakaratne, N. J., Burdick, J. W. & Roy, R. R. 2008. Training locomotor networks. Brain Res Rev, 57, 241–54.

Fortun, J., Puzis, R., Pearse, D. D., Gage, F. H. & Bunge, M. B. 2009. Muscle injection of AAV-NT3 promotes anatomical reorganization of CST axons and improves behavioral outcome following SCI. J Neurotrauma, 26, 941–53.

Friese, A., Kaltschmidt, J. A., Ladle, D. R., Sigrist, M., Jessell, T. M. & Arber, S. 2009. Gamma and alpha motor neurons distinguished by expression of transcription factor Err3. Proc Natl Acad Sci U S A, 106, 13588–93.

Gonzenbach, R. R., Gasser, P., Zorner, B., Hochreutener, E., Dietz, V. & Schwab, M. E. 2010. Nogo-A antibodies and training reduce muscle spasms in spinal cord-injured rats. Ann Neurol, 68, 48–57.

Han, Q., Ordaz, J. D., Liu, N. Richardson, Z., Wu, W., Xia, Y., Qu, W., Wang, Y., Dai, H., Zhang, Y. P., Shields, C. B., Smith, G. M. & Xu, X. M. 2019. Descending motor circuitry required for NT-3 mediated locomotor recovery after spinal cord injury in mice. Nat Commun, 10, 5815.

Hijikata, T., Wakisaka, H. & Yohro, T. 1992. Architectural design, fiber-type composition, and innervation of the rat rectus abdominis muscle. The Anatomical Record, 234, 500–512.

Jordan, L. M., Liu, J., Hedlund, P. B., Akay, T. & Pearson, K. G. 2008. Descending command systems for the initiation of locomotion in mammals. Brain Res Rev, 57, 183–91.

Kakanos, S. G. & Moon, L. D. F. 2019. Delayed peripheral treatment with neurotrophin-3 improves sensorimotor recovery after central nervous system injury. Neural Regeneration Research, 14, 1703–1704.

Kathe, C., Hutson, T. H., Mcmahon, S. B. & Moon, L. D. 2016a. Intramuscular Neurotrophin-3 normalizes low threshold spinal reflexes, reduces spasms and improves mobility after bilateral corticospinal tract injury in rats. Elife, 5.

Kathe, C., Hutson, T. H., Mcmahon, S. B. & Moon, L. D. 2016b. Intramuscular Neurotrophin-3 normalizes low threshold spinal reflexes, reduces spasms and improves mobility after bilateral corticospinal tract injury in rats. eLife, 5, e18146.

Kathe, C. & Moon, L. D. F. 2018. RNA sequencing dataset describing transcriptional changes in cervical dorsal root ganglia after bilateral pyramidotomy and forelimb intramuscular gene therapy with an adeno-associated viral vector encoding human neurotrophin-3. Data In Brief, 21, 377–385.

Krupka, A. J., Fischer, I. & Lemay, M. A. 2017. Transplants of Neurotrophin-Producing Autologous Fibroblasts Promote Recovery of Treadmill Stepping in the Acute, Sub-Chronic, and Chronic Spinal Cat. J Neurotrauma, 34, 1858–1872.

Langlet, C., Leblond, H. & Rossignol, S. 2005. Mid-lumbar segments are needed for the expression of locomotion in chronic spinal cats. J Neurophysiol, 93, 2474–88.

Lee-Kubli, C. A. G. & Calcutt, N. A. 2014. Altered rate-dependent depression of the spinal H-reflex as an indicator of spinal disinhibition in models of neuropathic pain. Pain, 155, 250–260.

Lee, S., Toda, T., Kiyama, H. & Yamashita, T. 2014. Weakened rate-dependent depression of Hoffmann’s reflex and increased motoneuron hyperactivity after motor cortical infarction in mice. Cell Death Dis, 5, e1007.

Maynard, F. M., Karunas, R. S. & Waring, W. P., 3RD 1990. Epidemiology of spasticity following traumatic spinal cord injury. Arch Phys Med Rehabil, 71, 566–9.

Mihai, G., Nout, Y. S., Tovar, C. A., Miller, B. A., Schmalbrock, P., Bresnahan, J. C. & Beattie, M. S. 2008. Longitudinal Comparison of Two Severities of Unilateral Cervical Spinal Cord Injury Using Magnetic Resonance Imaging in Rats. Journal of Neurotrauma, 25, 1–18.

Mills, C. D., Hains, B. C., Johnson, K. M. & Hulsebosch, C. E. 2001. Strain and model differences in behavioral outcomes after spinal cord injury in rat. J Neurotrauma, 18, 743–56.

Mohan, R., Tosolini, A. P. & Morris, R. 2014. Targeting the motor end plates in the mouse hindlimb gives access to a greater number of spinal cord motor neurons: an approach to maximize retrograde transport. Neuroscience, 274, 318–30.

Morris, R. & Whishaw, I. Q. 2016. A Proposal for a Rat Model of Spinal Cord Injury Featuring the Rubrospinal Tract and its Contributions to Locomotion and Skilled Hand Movement. Front Neurosci, 10, 5.

Mukherjee, A. & Chakravarty, A. 2010. Spasticity mechanisms - for the clinician. Front Neurol, 1, 149.

Musienko, P. E., Deliagina, T. G., Gerasimenko, Y. P., Orlovsky, G. N. & Zelenin, P. V. 2014. Limb and trunk mechanisms for balance control during locomotion in quadrupeds. The Journal of neuroscience: the official journal of the Society for Neuroscience, 34, 5704–5716.

Nishimura, Y., Onoe, H., Morichika, Y., Perfiliev, S., Tsukada, H. & Isa, T. 2007. Time-dependent central compensatory mechanisms of finger dexterity after spinal cord injury. Science, 318, 1150–5.

Ollivier-Lanvin, K., Fischer, I., Tom, V., Houle, J. D. & Lemay, M. A. 2015. Either brain-derived neurotrophic factor or neurotrophin-3 only neurotrophin-producing grafts promote locomotor recovery in untrained spinalized cats. Neurorehabil Neural Repair, 29, 90–100.

Pandyan, A. D., Gregoric, M., Barnes, M. P., Wood, D., Van Wijck, F., Burridge, J., Hermens, H. & Johnson, G. R. 2005. Spasticity: clinical perceptions, neurological realities and meaningful measurement. Disabil Rehabil, 27, 2–6.

Park, J. H., Kim, J. H., Oh, S.-K., Baek, S. R., Min, J., Kim, Y. W., Kim, S. T., Woo, C.-W. & Jeon, S. R. 2016. Analysis of equivalent parameters of two spinal cord injury devices: the New York University impactor versus the Infinite Horizon impactor. The Spine Journal, 16, 1392–1403.

Parkman, H. P., Rao, S. S., Reynolds, J. C., Schiller, L. R., Wald, A., Miner, P. B., Lembo, A. J., Gordon, J. M., Drossman, D. A., Waltzman, L., Stambler, N. & Cedarbaum, J. M. 2003. Neurotrophin-3 improves functional constipation. Am J Gastroenterol, 98, 1338–47.

Petruska, J. C., Kitay, B., Boyce, V. S., Kaspar, B. K., Pearse, D. D., Gage, F. H. & Mendell, L. M. 2010. Intramuscular AAV delivery of NT-3 alters synaptic transmission to motoneurons in adult rats. Eur J Neurosci, 32, 997–1005.

Pickett, W., Simpson, K., Walker, J. & Brison, R. J. 2003. Traumatic spinal cord injury in Ontario, Canada. J Trauma, 55, 1070–6.

Popovich, P. G., Tovar, C. A., Wei, P., Fisher, L., Jakeman, L. B. & Basso, D. M. 2012. A reassessment of a classic neuroprotective combination therapy for spinal cord injured rats: LPS/pregnenolone/indomethacin. Exp Neurol, 233, 677–85.

Reed, W. R., Shum-Siu, A., Onifer, S. M. & Magnuson, D. S. K. 2006. Inter-enlargement pathways in the ventrolateral funiculus of the adult rat spinal cord. Neuroscience, 142, 1195–1207.

Rossignol, S. & Frigon, A. 2011. Recovery of locomotion after spinal cord injury: some facts and mechanisms. Annu Rev Neurosci, 34, 413–40.

Ruder, L., Takeoka, A. & Arber, S. 2016. Long-Distance Descending Spinal Neurons Ensure Quadrupedal Locomotor Stability. Neuron, 92, 1063–1078.

Ryu, Y., Ogata, T., Nagao, M., Sawada, Y., Nishimura, R. & Fujita, N. 2021. Early escitalopram administration as a preemptive treatment strategy against spasticity after contusive spinal cord injury in rats. Sci Rep, 11, 7120.

Sahenk, Z. 2007. Pilot clinical trial of NT-3 in CMT1A patients. Prog Neurotherapeutics Neuropsychopharm, 2, 97–108.

Sahenk, Z., Galloway, G., Clark, K. R., Malik, V., Rodino-Klapac, L. R., Kaspar, B. K., Chen, L., Braganza, C., Montgomery, C. & Mendell, J. R. 2014. AAV1.NT-3 gene therapy for Charcot-Marie-Tooth neuropathy. Mol Ther, 22, 511–21.

Sahenk, Z., Nagaraja, H. N., Mccracken, B. S., King, W. M., Freimer, M. L., Cedarbaum, J. M. & Mendell, J. R. 2005. NT-3 promotes nerve regeneration and sensory improvement in CMT1A mouse models and in patients. Neurol, 65, 681–9.

Sangari, S. & Perez, M. A. 2019. Imbalanced Corticospinal and Reticulospinal Contributions to Spasticity in Humans with Spinal Cord Injury. J Neurosci, 39, 7872–7881.

Schucht, P., Raineteau, O., Schwab, M. E. & Fouad, K. 2002. Anatomical correlates of locomotor recovery following dorsal and ventral lesions of the rat spinal cord. Exp Neurol, 176, 143–53.

Singh, A., Murray, M. & Houle, J. D. 2011. A training paradigm to enhance motor recovery in contused rats: effects of staircase training. Neurorehabil Neural Repair, 25, 24–34.

Singh, A., Tetreault, L., Kalsi-Ryan, S., Nouri, A. & Fehlings, M. G. 2014. Global prevalence and incidence of traumatic spinal cord injury. Clin Epidemiol, 6, 309–31.

Skold, C., Levi, R. & Seiger, A. 1999. Spasticity after traumatic spinal cord injury: nature, severity, and location. Arch Phys Med Rehabil, 80, 1548–57.

Smith, C. C., Kissane, R. W. P. & Chakrabarty, S. 2018. Simultaneous Assessment of Homonymous and Heteronymous Monosynaptic Reflex Excitability in the Adult Rat. eNeuro, 5.

Smith, R. R., Shum-Siu, A., Baltzley, R., Bunger, M., Baldini, A., Burke, D. A. & Magnuson, D. S. 2006. Effects of swimming on functional recovery after incomplete spinal cord injury in rats. J Neurotrauma, 23, 908–19.

Sydney-Smith, J. D., Megaro, V., Spejo, A. B. & Moon, L. D. F. 2021a. An Adeno-Associated Viral vector encoding Neurotrophin 3 injected into affected forelimb muscles modestly improves sensorimotor function after contusive mid-cervical spinal cord injury. BioRxiv.

Sydney-Smith, J. D., Spejo, A. B., Warren, P. M. & Moon, L. D. F. 2021b. Peripherally delivered Adeno-associated viral vectors for spinal cord injury repair. Exp Neurol, 348, 113945.

Takeoka, A., Vollenweider, I., Courtine, G. & Arber, S. 2014. Muscle spindle feedback directs locomotor recovery and circuit reorganization after spinal cord injury. Cell, 159, 1626–39.

Talpalar, A. E., Bouvier, J., Borgius, L., Fortin, G., Pierani, A. & Kiehn, O. 2013. Dual-mode operation of neuronal networks involved in left-right alternation. Nature, 500, 85–8.

Tohyama, T., Kinoshita, M., Kobayashi, K., Isa, K., Watanabe, D., Kobayashi, K., Liu, M. & Isa, T. 2017. Contribution of propriospinal neurons to recovery of hand dexterity after corticospinal tract lesions in monkeys. Proc Natl Acad Sci U S A, 114, 604–609.

Wang, Y., Wu, W., Wu, X., Sun, Y., Zhang, Y. P., Deng, L. X., Walker, M. J., Qu, W., Chen, C., Liu, N. K., Han, Q., Dai, H., Shields, L. B., Shields, C. B., Sengelaub, D. R., Jones, K. J., Smith, G. M. & Xu, X. M. 2018. Remodeling of lumbar motor circuitry remote to a thoracic spinal cord injury promotes locomotor recovery. Elife, 7.

Watson, C., Paxinos, G., Kayalioglu, G. & Heise, C. 2009. Chapter 15 - Atlas of the Rat Spinal Cord. In: Watson, C., Paxinos, G. & Kayalioglu, G. (eds.) The Spinal Cord. San Diego: Academic Press.

Xu, N., Åkesson, E., Holmberg, L. & Sundström, E. 2015. A sensitive and reliable test instrument to assess swimming in rats with spinal cord injury. Behav Brain Res, 291, 172–183.

Yin, X., Yu, T., Chen, B., Xu, J., Chen, W., Qi, Y., Zhang, P., Li, Y., Kou, Y., Ma, Y., Han, N., Wan, P., Luo, Q., Zhu, D. & Jiang, B. 2019. Spatial Distribution of Motor Endplates and its Adaptive Change in Skeletal Muscle. Theranostics, 9, 734–746.

Zhou, L., Baumgartner, B. J., Hill-Felberg, S. J., Mcgowen, L. R. & Shine, H. D. 2003. Neurotrophin-3 expressed in situ induces axonal plasticity in the adult injured spinal cord. J Neurosci, 23, 1424–31.

Zorner, B., Bachmann, L. C., Filli, L., Kapitza, S., Gullo, M., Bolliger, M., Starkey, M. L., Rothlisberger, M., Gonzenbach, R. R. & Schwab, M. E. 2014. Chasing central nervous system plasticity: the brainstem’s contribution to locomotor recovery in rats with spinal cord injury. Brain, 137, 1716–32.

Zörner, B., Filli, L., Starkey, M. L., Gonzenbach, R., Kasper, H., RöThlisberger, M., Bolliger, M. & Schwab, M. E. 2010. Profiling locomotor recovery: comprehensive quantification of impairments after CNS damage in rodents. Nat Methods, 7, 701–8.

