## Supplementary figures 1-3 for "Delayed viral vector mediated delivery of neurotrophin-3 improves skilled hindlimb function and stability after thoracic contusion in rats"

Supplementary Figure 1: Thoracic contusion injury causes bilateral effects on hindlimb function.

Supplementary Figure 2: Thoracic contusion permanently increases hindlimb miss-stepping in skilled tasks

Supplementary Figure 3: H reflex parameters did not alter in unexcluded animals following thoracic contusion.

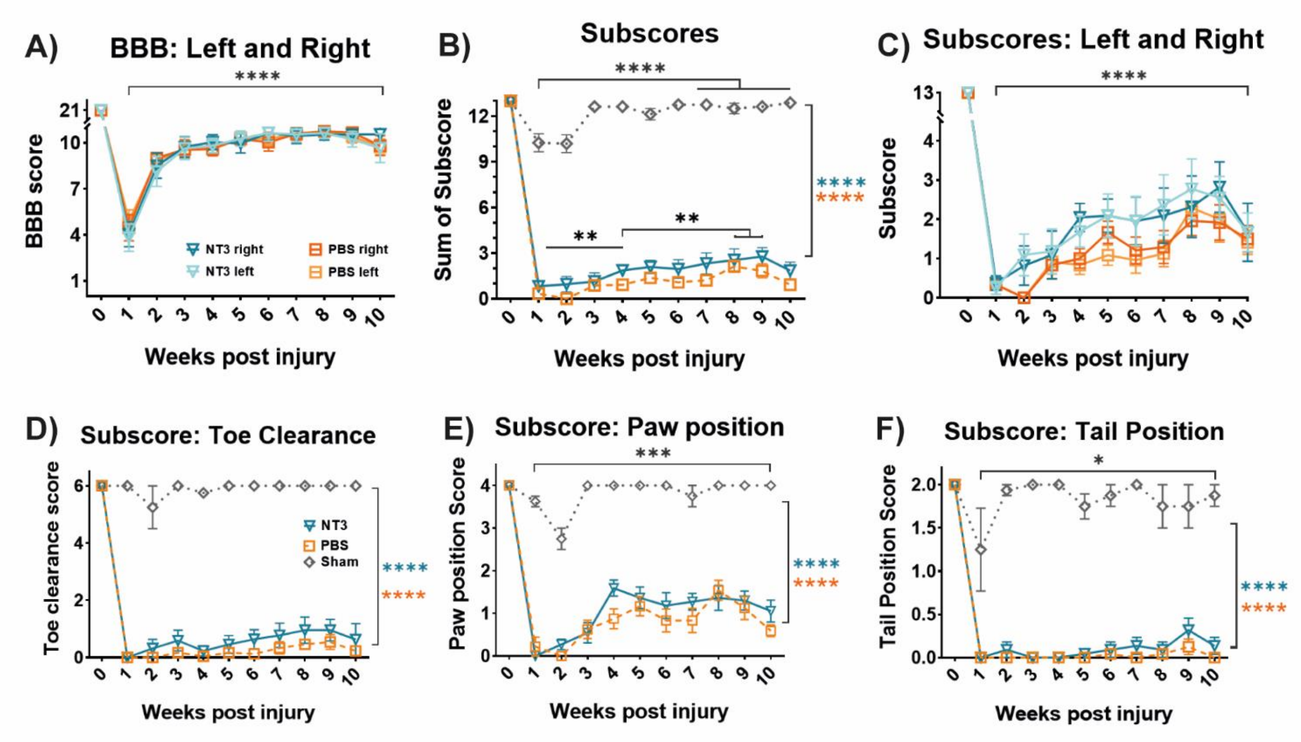


**Supplementary Figure 1:** Thoracic contusion injury causes bilateral effects on hindlimb function. A) BBB scores showed no functional difference between the injury induced deficits of the left and right hindlimbs of either the NT3 or control groups, indicating that the contusion was functionally bilateral. Left = light colours, right = dark colours. B) BBB subscores were reduced after injury, and not improved by NT3 treatment compared to controls (effect of group, linear model, $F(2, 24)=178$, $p<0.0001$, post hoc LSD comparison, NT3 vs Naïve $p<0.0001$, PBS vs Naïve $p<0.0001$, NT3 vs PBS $p=0.089$). C) Deficits in BBB subscore were comparable between left and right hindlimbs in all injured animals. Left = light colours, right = dark colours. D-F) Component sections of the BBB were analysed showing D) average toe clearance in which the NT3 group had a trend towards higher average toe clearance, no evidence for an effect of NT3 was detected (effect of time x injury group, linear model, $F(18,211)=0.58$, $p=0.91$. E) Average paw position was abnormal after injury and NT3 treatment caused little overall improvement. F) Tail positioning during locomotion was altered after injury. However, NT3 did not cause a recovery in this specific function (effect of time x group, linear model, $F(18,215)=2.05$, $p=0.0085$, post hoc LSD, NT3 vs PBS at all timepoints $p>0.19$). In all panels: NT3 = blue triangle, controls = orange square, sham = black diamond, and wpi = week post injury. * = $P<0.05$, ** = $P<0.01$, *** = $P<0.001$, and **** = $P<0.0001$. If no post-hoc result is shown, comparison was not-significant. Values represent mean±SEM.

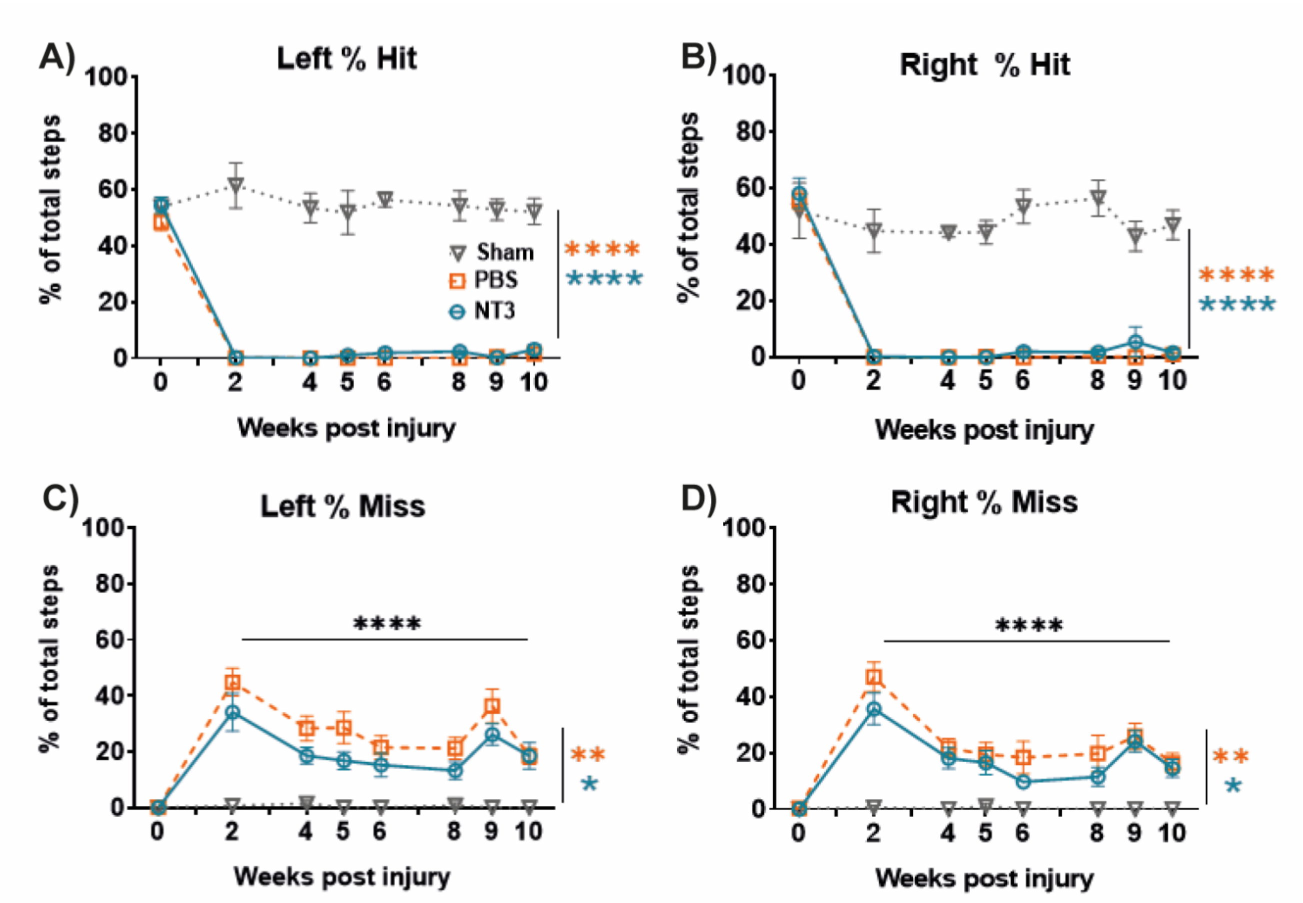


**Supplementary Figure 2:** Thoracic contusion permanently increases hindlimb miss-stepping in skilled tasks. The thoracic contusion caused a permanent decrease in the number of error-free steps in the A) left and B) right hindlimbs (effect of group, linear model, left = F(2,23)=910, P<0.0001, post hoc LSD for NT3 vs Sham P<0.0001, PBS vs Sham P<0.0001, right = F(2,24)=693.6, P<0.0001, post hoc Fisher's LSD for NT3 vs Sham P<0.0001, PBS vs Sham P<0.0001 and NT3 vs PBS p=0.17) with no increase shown in the NT3 group compared to injured control (NT3 vs PBS, left = p=0.40, right = p=0.17). C-D) Further, the contusion injury caused a substantive increase in the number of bilateral miss-steps performed by all animals (effect of group, linear model, left = F(2,24)=9.95, P=0.0007, post hoc LSD for NT3 vs Sham P=0.0053, PBS vs Sham P=0.0002, NT3 vs PBS p=0.079, right = F(2,24)=5.68, P=0.0095, post hoc LSD for NT3 vs Sham P=0.017, PBS vs Sham P=0.0025, NT3 vs PBS p=0.30) which was similarly not recovered following NT3 treatment. Baseline recordings were not used as a covariate but were excluded for tests of effects and interactions. For NT3 n=11 for PBS n=11, for sham n=4. In all panels: NT3 = blue triangle, controls = orange square, sham = black diamond, and wpi = week post injury. * = P<0.05, ** = P<0.01, *** = P<0.001, and **** = P<0.0001. If no post-hoc result is shown, comparison was not-significant. Values represent mean±SEM.

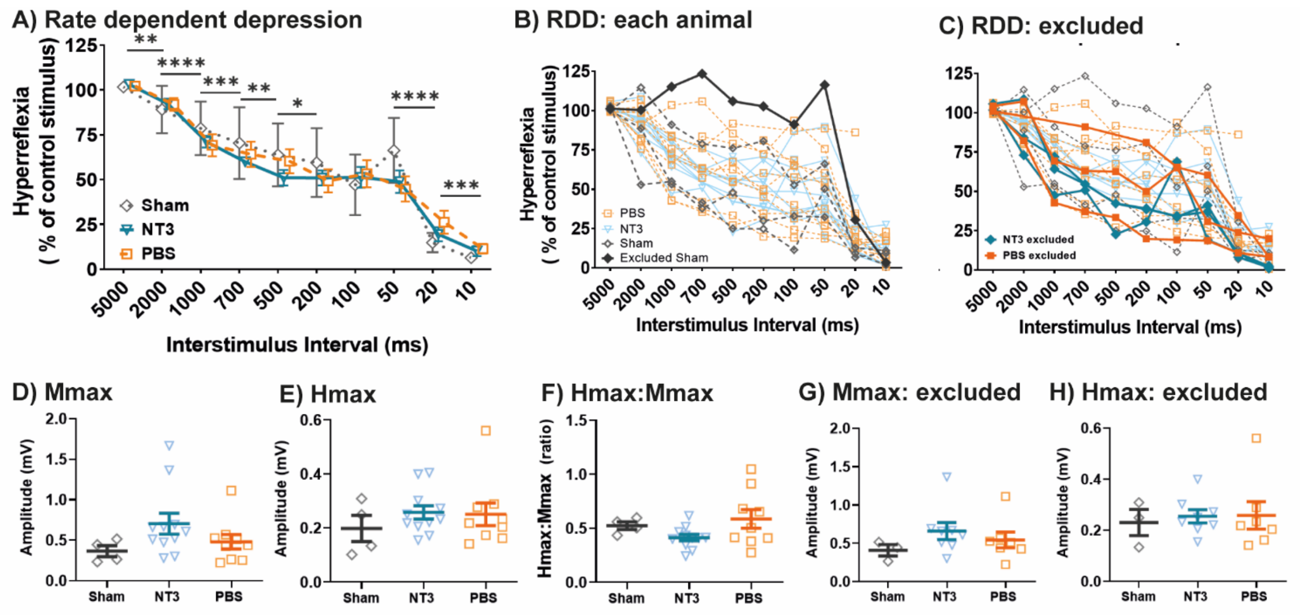


**Supplementary Figure 3:** H reflex parameters did not alter in unexcluded animals following thoracic contusion. A) H wave of all animals showing RDD was present in comparable amounts in both injured and sham animals (effect of group, linear model, F(2,22)=0.14, P=0.87), with no treatment interaction present (effect of group x interstimulus interval, linear model, F(18,185)=1.09, P=0.42). B) RDD for every animal prior to exclusion. C) RDD of animals excluded from analysis showing similar trends to the group data. D) The maximum amplitude of the M wave was comparable between all contused and sham animals (one way ANOVA, F(2,21)=1.86, p=0.18). E) The maximum amplitude of the H wave was comparable between all contused and sham animals (one way ANOVA, F(2,21)=0.50, p=0. 61). F) The Hmax:Mmax ratio, an indicator of reflex excitability, was comparable between all animals of the injured and sham groups (One way ANOVA, F(2,21)=2.41, p=0.11). G) The maximum amplitude of the M wave was comparable between contused and sham animals following exclusions (one way ANOVA, F(2,15)=0.94, p=0.41). H) The maximum amplitude of the H wave was comparable between contused and sham animals following exclusions (one way ANOVA, F(2,15)=0.076, p=0.93).
